# α-Parvin regulation of cell re-arrangement is critical for ureteric bud branching morphogenesis

**DOI:** 10.64898/2025.12.04.692378

**Authors:** Xinyu Dong, Fabian Bock, Ali Hashmi, Nada Bulus, Glenda Mernaugh, Gema Bolas, Shensen Li, Wanying Zhu, Meiling Melzer, Kyle Brown, Colton Miller, Olga Viquéz, Eloi Montañez, Ambra Pozzi, Sara A. Wickström, Roy Zent

## Abstract

All branched tubular structures, including the kidney collecting system, are formed by branching morphogenesis, a process that includes tip branching and trunk narrowing. Tight control of cell movement and rearrangements is a prerequisite for branching morphogenesis. The role of integrin-associated adhesion proteins in coordinating actin dynamics and cell rearrangements during branching morphogenesis is poorly understood. Here we used 3D live imaging of mouse ureteric bud branching to show that α-parvin, a component of the integrin binding ILK-PINCH-Parvin (IPP) complex, regulates tip branching and tubule thinning by inhibiting excessive cell adhesion and actin polymerization. Mechanistically, α-parvin promotes actin turnover by inhibiting activation of the small GTPases RhoA and Cdc42, which in turn enhances the severing function of the actin regulatory protein, cofilin. These results underscore the importance of adhesion protein-regulated actin dynamics in the critical process of cell rearrangement, which is required for branching morphogenesis.

**Teaser:** Scaffold protein α-parvin limits cell adhesion and actin polymerization, enabling cell rearrangements that drive kidney branching

## Introduction

Branching morphogenesis underlies the formation of all branched tubular structures like the developing lung, mammary gland, and kidney (1). This process requires tightly coordinated cell movement and rearrangements (2). While extensive information exists on how growth factor-dependent signaling regulates branching morphogenesis (3), significantly less is known about the role of cell-extracellular matrix (ECM) interactions and actin dynamics during this process, especially in kidney development (4, 5).

The kidney collecting duct system is formed from the ureteric bud (UB), a single-layered epithelial tube with a lumen that grows out of the nephric duct into the metanephric mesenchyme (6, 7). The mouse UB tips undergo multiple cycles of branching, and the newly formed tubules undergo extensive thinning and elongation giving rise to the collecting ducts (CD) (8–10). The mechanisms and molecules that drive this complex morphogenic process are still poorly understood.

Cell-ECM interactions mediated by integrins are critical for UB branching morphogenesis. Integrins are transmembrane α/β heterodimers that link extracellular cues to intracellular processes like signal transduction and cytoskeletal remodeling, which allow them to mediate cell functions like adhesion, migration and cytoskeletal organization. Integrin function is mediated by the extracellular domain binding to ECM and the cytoplasmic tails recruiting scaffold proteins (11) including the ILK/Pinch/parvin complex (12, 13). ILK (14), the best studied component of this complex, promotes integrin dependent adhesion and migration (15, 16) and reduces actomyosin contractility (17, 18). Parvins (14, 19) are a three member family consisting of the ubiquitously expressed α-parvin; β-parvin, which is primarily expressed in heart and skeletal muscle; and hematopoietic γ-parvin (20). In addition to binding to ILK, parvins interact with filamentous actin (F-actin) (21) and actin-regulatory proteins including regulators of Rho GTPases (22, 23). They are thought to have similar functions to ILK (24, 25). Germ line deletion of α-parvin in mice led to kidney agenesis (26), however the role of α-parvin in UB branching morphogenesis is still unknown.

In this study, we used state-of-the-art 3D live-organ imaging to track individual cell behaviors in whole kidneys, to show that α-parvin promotes coordinated cell rearrangements that drive both UB branching and tubule thinning. Unexpectedly, loss of α-parvin caused increased cell adhesion, aberrant migration, and excessive F-actin formation, which was unlike the phenotype seen when β1 integrins or ILK were deleted in the developing UB. Mechanistically, α-parvin promoted actin turnover by inhibiting RhoA and Cdc42 activation, which in turn enhanced the actin-severing function of cofilin. These findings uncover unexpected divergent roles between α-parvin and ILK during epithelial morphogenesis and provide insights into how various components within adhesion complexes differentially regulate actin dynamics during tissue morphogenesis. It also underscores the general importance of regulated actin remodeling in branching morphogenesis, a process fundamental to the development of multiple organ systems including the kidney, lung, mammary gland, and pancreas.

## Results

### Deleting α-parvin in the UB causes end stage kidney failure

Deleting α-parvin resulted in renal agenesis, which suggested it is expressed early in the developing kidney (26). We verified that α-parvin is present in both the stalk and tips of the developing UB at E12.5 (Fig S1A), as well as in tubules and glomeruli (derived from the MM) of newborn mice (Fig S1B). To identify the role of α-parvin specifically in the UB, we selectively deleted it by crossing floxed α-parvin mice (α-pv^f/f^) with HoxB7^Cre^ mice, which express cre recombinase in the Wolffian duct and UB starting at E10.5 (27). We confirmed deletion in the cytokeratin positive CDs (which are derived from the UB) in the postnatal day 1 Hoxb7:α-pv^f/f^ mice (Fig 1A) and the developing UB at E12.5 (Fig S1A). The Hoxb7:α-pv^f/f^ mice were born in the predicted Mendelian ratio (Fig S1C); however, most of them died by 6 months of age (Fig 1B). Kidneys isolated from the Hoxb7:α-pv^f/f^ mice just prior to death were hypoplastic and dysmorphic and there were large areas of inflammation with tissue destruction in both the cortex and medulla making it difficult to identify mechanisms for the phenotype (Fig 1C). We therefore examined 2-month-old mice, where the tissue destruction was less severe. At this age Hoxb7:α-pv^f/f^ mice already had severe renal dysfunction with serum blood urea nitrogen levels significantly higher than in α-pv^f/f^ mice (Fig S1C). The kidneys were significantly smaller than the α-pv^f/f^ kidneys and the ureters inserted into the bladder with no evidence of ureteric obstruction (Fig 1D). The papillae in these mice were hypoplastic (Fig 1E) and both the cortical and medullary CDs were disordered and dilated (Fig 1E). Inflammatory cell infiltration was present (Fig 1E) and fibrosis, quantified using Sirius red staining, was significantly increased (Fig 1F-G). There were decreased AQP2 positive tubules in the Hoxb7:α-pv^f/f^ kidneys (Fig 1H) suggesting CD development defects. Thus, deleting α-parvin results in an end stage kidney characterized by a loss of kidney structures, a hypoplastic collecting system and severe fibrosis and inflammation in both the kidney cortex and medulla.

**Figure 1.**
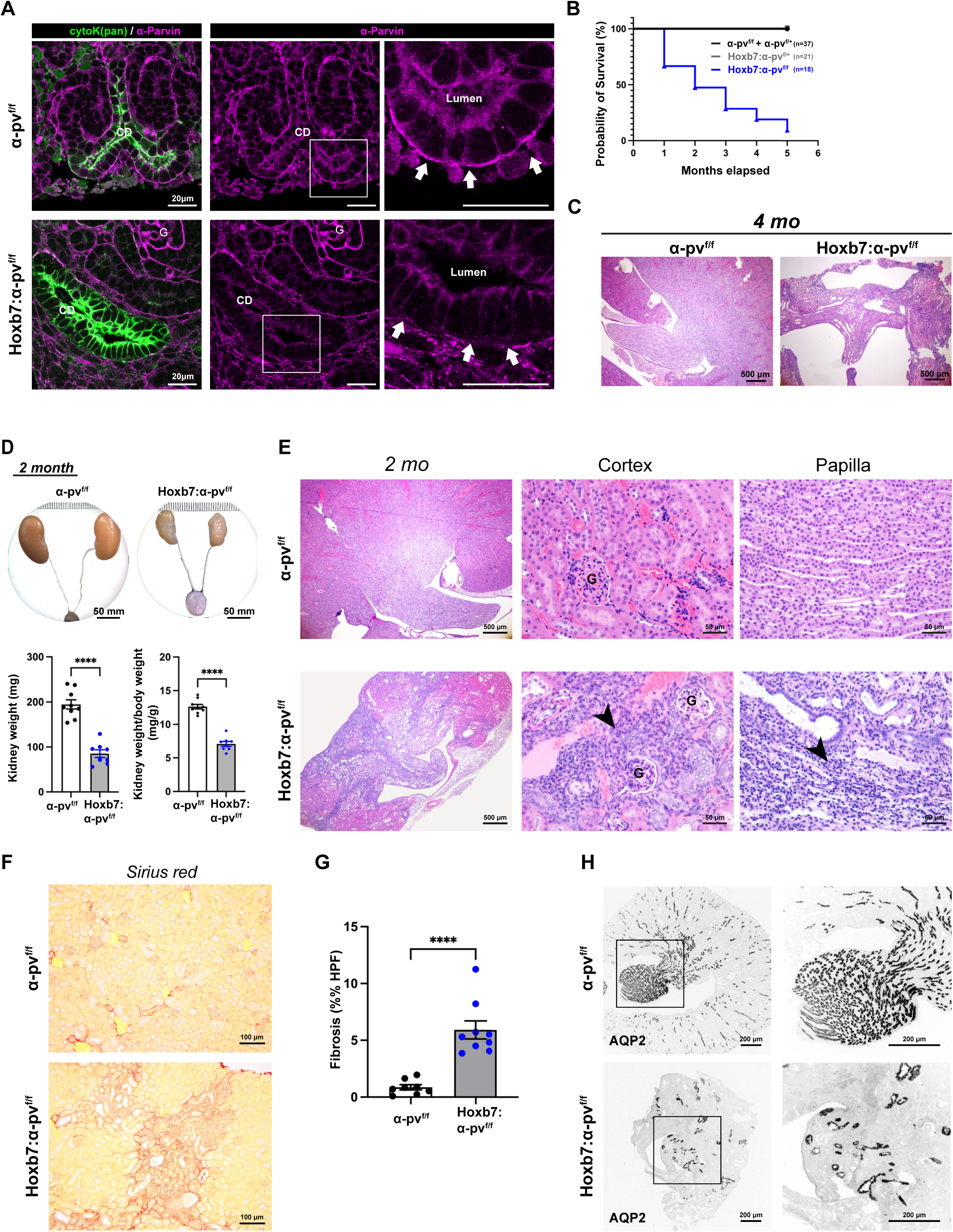
Deleting α-parvin in the UB results in a branching morphogenesis defect and kidney failure. **(A)** Representative images of α-parvin immunostaining performed on frozen kidney sections of newborn α-pv^f/f^ and Hoxb7:α-pv^f/f^ kidneys. The collecting ducts (CD) were labeled with dolichus biflorus agglutinin (DBA) and α-parvin (magenta). α-parvin was expressed on the baso-lateral sides (arrows) of the CD cells of α-pv^f/f^ mice. There was a decrease in α-parvin staining in the CDs (arrows) but not the glomeruli (G) in the Hoxb7:α-pv^f/f^ kidneys. 3 mice per group were analyzed. **(B)** A Kaplan-Meier curve analysis revealed that 50% of Hoxb7:α-pv^f/f^ mice die within 2 months of birth and they were all dead by 6 months. There was no mortality in α-pv^f/f^ or Hoxb7:α-pv^f/+^ mice over 6 months (p < 0.0001). **(C)** Hematoxylin and eosin (H&E) staining of kidney sections from 4-month-old α-pv^f/f^ and Hoxb7:α-pv^f/f^ mice demonstrating destruction of the cortex and severe hypoplasia and dysplasia of the medulla. n = 3 mice per group. **(D)** Representative image of dissected kidneys with ureters and bladder from 2-month-old α-pv^f/f^ and Hoxb7:α-pv^f/f^ mice. Ureters in Hoxb7:α-pv^f/f^ mice insert normally into the bladder without signs of ureteric dilation or obstruction (upper panel). Total kidney weight (bottom right) and kidney-to-body weight ratios (bottom left) at 2 months of age revealed significantly smaller kidneys in Hoxb7:α-pv^f/f^ mice compared to α-pv^f/f^ littermates ((α-pv^f/f^, n = 9; Hoxb7:α-pv^f/f^, n = 8)). Data are shown as mean ± SEM. ****p < 0.0001; 2 tailed t-test. **(E)** H&E staining of kidneys from 2-month-old α-pv^f/f^ and Hoxb7:α-pv^f/f^. Low magnification (left panels) demonstrated fibrosis and hypoplasia/dysplasia of the Hoxb7:α-pv^f/f^ kidneys. Higher magnification of the cortex with the inclusion of glomeruli (G) (center panel) and medulla (right panel) demonstrated disorganized CDs and excessive inflammatory cells (arrowheads) in Hoxb7:α-pv^f/f^ kidneys. n = 8 mice per group. **(F-G)** Picrosirius red staining of kidneys from 2-month-old α-pv^f/f^ and Hoxb7:α-pv^f/f^ revealed more fibrosis in Hoxb7:α-pv^f/f^ mice compared to α-pv^f/f^ mice (F). The area of 5 high-power fields per sample of red stained area was quantified using ImageJ. The data are expressed as mean ± SEM. n = 8 mice per group (G), ****p < 0.0001; 2 tailed t-test. **(H)** Representative immunostaining of kidneys from 2-month-old α-pv^f/f^ and Hoxb7:α-pv^f/f^ mice for AQP2 revealed less CDs in Hoxb7:α-pv^f/f^ kidneys. n = 3 mice per group were analyzed.

### Deleting α-parvin in the UB causes a branching morphogenesis defect

To define whether the hypoplastic papilla was caused by a UB branching morphogenesis defect, we dissected embryonic kidneys at E10.5 (soon after the initiation of UB development)(8) and visualized the developing UB with E-cadherin and the developing MM with Six2. The UB in the α-pv^f/f^ mice invaded the MM and an initial branching step was observed at E10.5. This process was delayed in the Hoxb7:α-pv^f/f^ mice where, although the UB invaded the MM, it did not branch (Fig 2A). This branching defect was confirmed in E12.5 kidneys, where the α-pv^f/f^ mice had approximately twice the number of branch tips compared to the Hoxb7:α-pv^f/f^ UBs (Fig 2A). The more severe branching defect was also seen in Hoxb7:α-pv^f/f^ E12.5 kidneys cultured on transwells for 16 hours, which is sufficient time for at least one round of branching (28) (Fig. 2A). Histological analysis of Hoxb7:α-pv^f/f^ embryonic kidneys at E15.5 revealed hypoplastic kidneys with a notable reduction in MM volume and UB branching compared to α-pv^f/f^ controls (Fig 2A). A similar but more pronounced phenotype was seen at E18.5, where the mutant kidneys were markedly smaller with a hypoplastic papilla, medulla, and cortex (Fig 2B and C). These kidneys did not have any evidence of ureteric obstruction (Fig S2A). The typical monolayered epithelial cell CD structure was disrupted, and cells appeared to pile up on each other within the CD lumens, resulting in enlarged tubules in the Hoxb7:α-pv^f/f^ mice when compared to the α-pv^f/f^ mice (Fig 2B and C). When we quantified the size of the CDs by measuring their diameters in kidney cross sections, the average diameter in Hoxb7:α-pv^f/f^ kidneys was 30.55 ± 9.3 μm compared to 20.91 ± 10.51 in the α-pv^f/f^ kidneys and there was a right-shift of the distribution of tubular size with an increased diameter in the mutant mice (Fig 2D).

**Figure 2.**
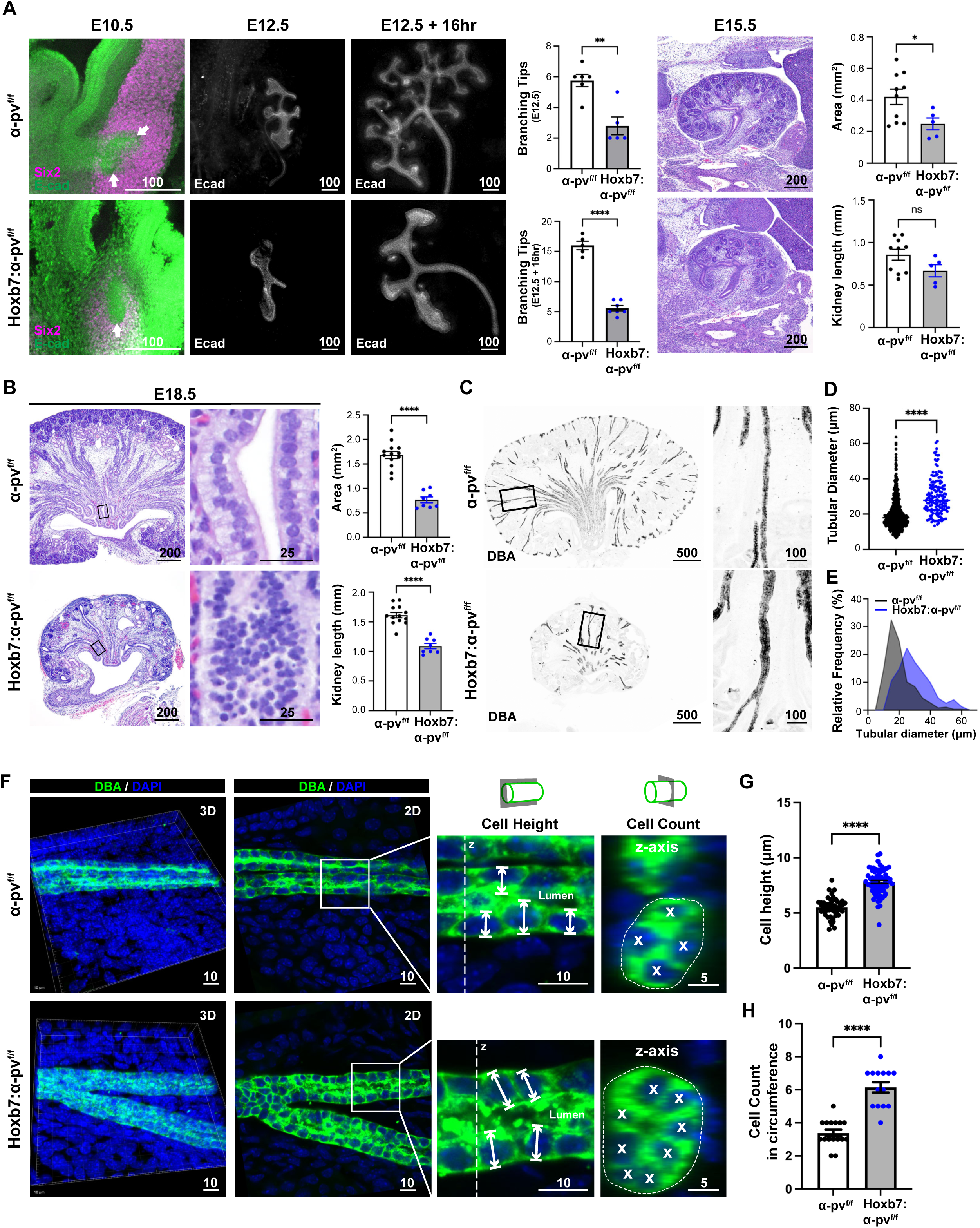
Deleting α-parvin in the UB causes a branching morphogenesis and tubule thinning defect. **(A)** Kidneys were isolated from embryos of α-pv^f/f^ and Hoxb7:α-pv^f/f^ mice at E10.5 (left), E12.5 (center), or isolated at E12.5 kidneys and cultured ex vivo for 16 hours (right). Whole mounts of E10.5 kidneys were stained with E-cadherin (indicating UB) and Six2 (indicating the MM). Imaging with confocal microscopy showed invasion (arrows) but delayed branching of the UB within the MM of the Hoxb7:α-pv^f/f^ mice. E12.5 kidneys and E12.5 kidneys cultured ex vivo (E12.5 + 16hr) stained with E-cadherin showed a branching defect in the Hoxb7:α-pv^f/f^ mice compared to the α-pv^f/f^ mice. Quantification showed significantly fewer branches in the Hoxb7:α-pv^f/f^ kidneys compared to the α-pv^f/f^ mice at both E12.5 (α-pv^f/f^, n=6; Hoxb7:α-pv^f/f^, n=5) and after 16 hours of ex vivo culture (α-pv^f/f^, n=5; Hoxb7:α-pv^f/f^, n=6). E15.5 α-pv^f/f^ and Hoxb7:α-pv^f/f^ stained with H&E demonstrated a branching defect and decreased MM development in the mutant mice. Hoxb7:α-pv^f/f^ kidneys were smaller in total area (upper panel) but not length (lower panel) compared to α-pv^f/f^ kidneys (α-pv^f/f^, n=10; Hoxb7:α-pv^f/f^, n=5). Data are shown as mean ± SEM. *p < 0.05; 2 tailed t-test. **(B)** E18.5 α-pv^f/f^ and Hoxb7:α-pv^f/f^ stained with H&E demonstrated a branching defect and decreased MM induction in the mutant mice. High power views demonstrate a disorganized and hypercellular CD in the tubules of the Hoxb7:α-pv^f/f^ mice. Hoxb7:α-pv^f/f^ kidneys were smaller in both total area (upper panel) and length (lower panel) than α-pv^f/f^ kidneys (α-pv^f/f^, n=13; Hoxb7:α-pv^f/f^, n=8). Data are shown as mean ± SEM. ****p < 0.0001; 2 tailed t-test. **(C-E)** Frozen sections of kidneys from newborn α-pv^f/f^ and Hoxb7:α-pv^f/f^ labelled with DBA show widened CDs in Hoxb7:α-pv^f/f^ kidney (C). Tubule diameters were measured as described in the methods. CD tubule widths are shown as mean ± SEM (D) and distribution plots (E). Tubules were pooled from 3 individual mice per genotype, ****p < 0.0001; 2 tailed t-test. **(F-H)** Thick frozen kidney sections of neonatal α-pv^f/f^ and Hoxb7:α-pv^f/f^ mice were immunolabelled with DBA and DAPI. They were imaged by confocal microscopy in 3D and then rotated to be horizontal prior to tubular shape analysis. Tubules were noted to only contain a single cell. The height (x-y plane) and number (z plane) of cells were counted in the tubules as described in the methods. Quantifications of cell heights (G) and cell number (H) were presented as mean ± SEM. Measurements were accumulated over 3 individual mice per genotype with at least 5 tubules analyzed, ****p < 0.0001; 2 tailed t-test.

To determine whether this expanded tubular size was due to increased cellular proliferation, we labelled the CD with Ki67. We found a significant decrease in cellular proliferation within the developing Hoxb7:α-pv^f/f^ CDs that were identified by cytokeratin staining (Fig S2C), which suggests that CD hyperproliferation was not the cause of the luminal hypercellularity. To investigate whether the widened CDs in the Hoxb7:α-pv^f/f^ kidneys were due to abnormal multilayering of the epithelium, we performed 3D confocal imaging of these tubules in thick frozen sections. The orthogonal views (z-axis) (Fig 2E) revealed that the tubules were a monolayer, however, the cells were significantly taller (Fig 2E, 2F) and there was a significant increase in the number of cells per cross section (Fig 2G) in the Hoxb7:α-pv^f/f^ kidneys when compared to the α-pv^f/f^ kidneys. These data suggest that the widening of the tubules was due to a morphological defect of the CD cells. Thus, deleting α-parvin in the UB appears to cause abnormal epithelial cell organization resulting in the inability to branch and a failure in establishing normal tubular geometry.

### α-parvin regulates CD cell neighbor exchange during UB branching

To determine the cellular mechanisms underlying the defective branching morphogenesis and tubule thinning of the UB the Hoxb7:α-pv^f/f^ mice, we used mTmG mice in which cells initially express the membrane-targeted red fluorescent protein “mT”, but upon Cre-mediated recombination, switches to express the membrane-targeted GFP “mG” (29). We crossed mTmG mice with Hoxb7:α-pv^f/f^ transgenic mice and performed volumetric confocal imaging at regular intervals. This allowed us to identify and track individual UB cells over an extended period within ex vivo cultured kidneys utilizing a methodology adopted from salivary gland (30) and embryonic kidney cultures (31). We isolated kidneys from these mice at E12.5, grew them on transwells (30, 31) (Fig. 3A)(movies 1-2) and utilized 3D confocal microscopy to capture branching morphogenesis at 10-minute intervals for a total of 22 hours. As expected, the UB tip in the α-pv^f/f^ kidney first enlarged to form an ampulla and then underwent remodeling to form two tips (Fig 3B, top panel). In contrast, the UB from Hoxb7:α-pv^f/f^ mice showed delayed ampulla enlargement and decreased trunk remodeling resulting in wide unbranched tubules (Fig 3B, bottom panel). To define the dynamics of tubule remodeling in greater detail, we examined trunk narrowing using orthogonal views of the time-lapse z-stacks (Fig 3C orange line and movies 3-4). The tubules were aligned with the x-axis and the cross-sectional area was measured over time. The elongating tubules in the α-pv^f/f^ mice narrowed 50% over time, while the tubule diameter in the Hoxb7:α-pv^f/f^ mice did not change over the same time (Fig 3C-3D).

**Figure 3.**
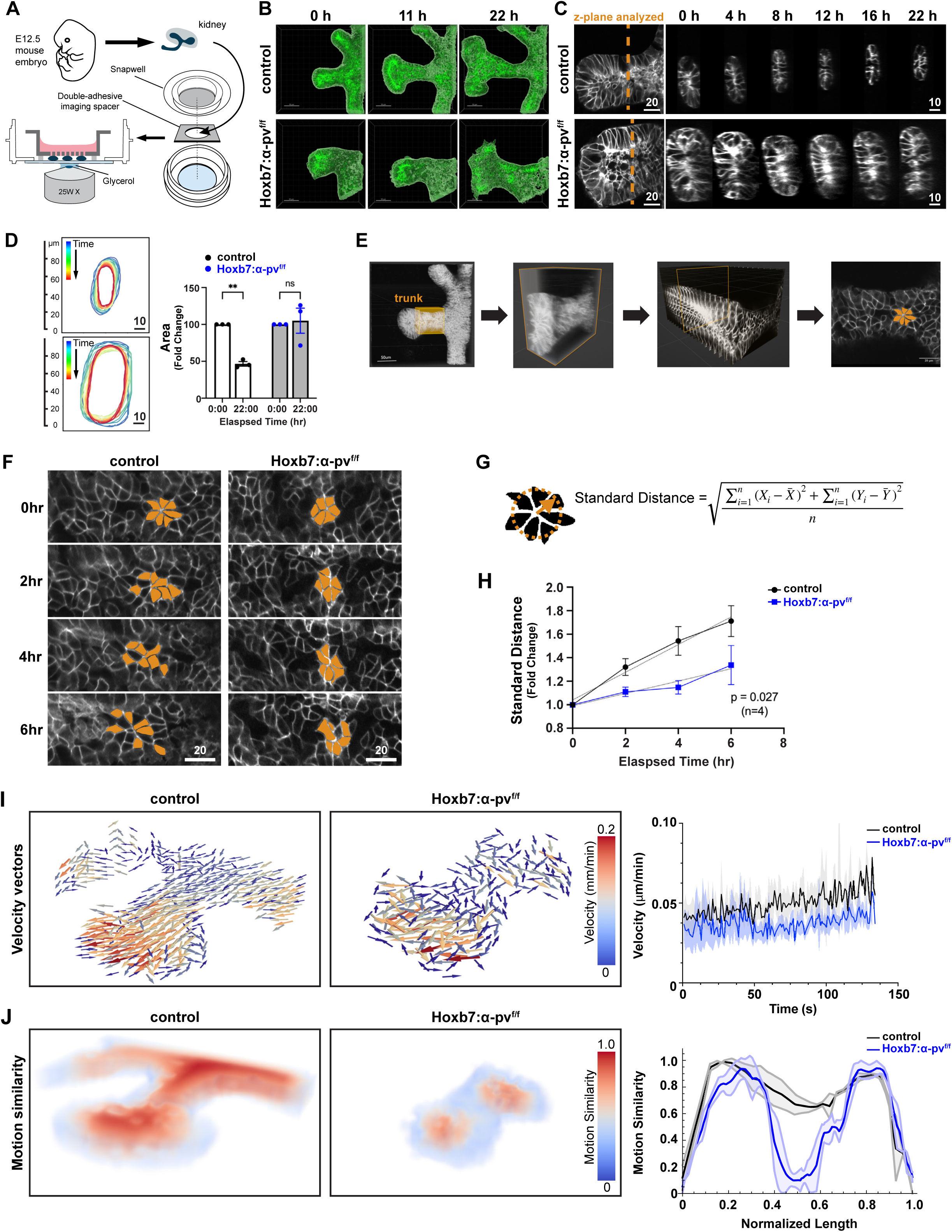
CD thinning defect in α-parvin null mice is due to abnormal cell rearrangements. **(A)** Schematic diagram of sample preparation for live imaging of wholemount mouse embryonic kidney. **(B)** E11.5 kidneys isolated from controls (control kidneys were Hoxb7:mTmG^f/f^ unless otherwise noted.) and Hoxb7:mTmG^f/f^:α-pv^f/f^ mice were cultured ex vivo. Confocal image stacks obtained at different time points demonstrate a branching and thinning defect in the Hoxb7:mTmG^f/f^:α-pv^f/f^ mice. **(C-D)** Time-lapse confocal microscopy images of isolated E11.5 UBs from control and Hoxb7:mTmG^f/f^:α-pv^f/f^ mice were taken over the time points shown and subjected to 3D reconstruction The tubules were rotated for cross-sectional analysis as depicted by an orange line (B). The UB surface was outlined in Imaris 10.0 to allow 3D rendering of the structure and the progression of stalk narrowing over time is depicted graphically against elapsed time (C). The relative change in cross-sectional area control and Hoxb7:mTmG^f/f^:α-pv^f/f^ mice over 22 hours is compared. UB Trunks from three individual kidneys per genotype were followed (mean ± SEM), **p < 0.01; Two-way ANOVA followed by Tukey’s test (D). (E) Workflow illustrating how the UB trunk region was selected for analysis. A defined region within the trunk (orange box) was extracted from z-stack confocal images and rendered in 3D. A single plane from this region, in which cell boundaries were visible via membranous GFP, was used for analysis of cell rearrangement. **(F-H)** Representative time-lapse images showing the dispersal of a rosette-like cell cluster in control and Hoxb7:mTmG^f/f^:α-pv^f/f^ collecting ducts (CDs) over 6 hours. Cells within the cluster were pseudocolored (orange) to visualize cell rearrangement over time. (F). The standard distance as defined in the methods was used to quantify the extent of cell dispersion during rearrangement (G). Standard distance for the cell clusters were plotted against elapsed time with a linear regression fit applied to the data. The p-value indicates a comparison between the slope of the two linear regression fits using the analysis of covariance (ancova). Four independent UB trunks per genotype were analyzed, with a minimum of 4 cell clusters from each trunk (mean ± SEM) (H). **(I and J)** Particle intensity velocity (PIV) analysis of the 3D live-images. (I) Representative images of velocity vector magnitude and orientation (left) and quantification of velocity (right) from live imaging of control and Hoxb7:mTmG^f/f^:α-pv^f/f^ tissue. Note reduced velocity and lack of global directionality of vectors (n= 5 (control), 4 (Hoxb7:mTmG^f/f^:α-pv^f/f^)). (J) Representative images of motion similarity heatmaps (left) and quantification of motion coherence (right) from live imaging of control and Hoxb7:α-pv^f/f^ tissue. Three equidistant parallel lines were drawn across the length of the tissue to determine the average and standard deviation of motion similarity. Note lack of persistent and coordinated global motion across the tissue in Hoxb7:α-pv^f/f^ relative to control n= 5 (control), 4 (Hoxb7:α-pv^f/f^) The full PIV analysis results are shown in supplementary movies S1 (velocity vector magnitude) and S2 (motion similarity heatmaps).

Tubular elongation and narrowing occur either by oriented cell division, as found in in late CD development (9), or by luminal mitosis like in UB branching tips (9, 32). To define which of these mechanisms occurred in the early elongating UB trunk, we followed dividing cells over time from our time-lapse imaging of the newly formed E12.5 trunks. We found that cell division was not oriented with the longitudinal axis of the tubule in either α-pv^f/f^ or Hoxb7:α-pv^f/f^ UB trunks (Fig S3A-C). However, we did observe that the dividing cells underwent luminal mitosis (32), a process where cells delaminate from the epithelium and divide within the lumen, in both α-pv^f/f^ and Hoxb7:α-pv^f/f^ CDs, as occurs in UB tips (Fig S3 D, E). Thus, normal UB trunk elongation occurs by luminal mitosis and this process is not impaired in Hoxb7:α-pv^f/f^ UB trunks. Furthermore, these results suggest that cell re-arrangement plays a role during the thinning process.

Cell re-arrangement, a conserved feature of tubular epithelial morphogenesis(33, 34), is required for UB development. This process is characterized by the formation and dispersal of multicellular rosette structures (35), where cells rapidly exchange their neighbors by performing controlled epithelial locomotion (36). To measure the cell-rearrangement, we marked multicellular rosettes within the tubule trunk (Fig 3E) and tracked the position of the individual cells in the rosettes over time (Fig 3F and movies 5-6). Rosettes were defined as a configuration of at least five cells that shared a vertex and spatial cell dispersal was measured and quantified (See Methods, Fig. 3G and H). There was a significant decrease in cell motility and neighbor exchange in Hoxb7:α-pv^f/f^ mice over 6 hr (Figure 3H).

The reduced neighbor exchanges suggested that the α-parvin-deficient tissue is in a less fluid-like state than the control tissue (37). To infer the mechanical phase state of the tissue, we quantified coarse-grained velocity fields across the tissue (38). Indeed, the average magnitude of the velocity field indicated slower motility of Hoxb7:α-pv^f/f^ tissues compared to α-pv^f/f^ mice (Fig 3I and movies 7-8). Probing the local directions and alignment of velocity vectors over time (motion similarity; Fig 3I and movies 9-10) further revealed coordinated cellular motion in α-pv^f/f^ that persisted over time, characteristic of fluid-like behavior. In contrast, Hoxb7:α-pv^f/f^ tissues did not maintain persistent, coordinated motion across the length of the tissue, characteristic of a jammed or solid-like state and consistent with the slower growth-rate of the tissues (39) (Fig 3I, J). Taken together these results indicate a decrease in neighbor exchanges and persistent motion of the Hoxb7:α-pv^f/f^ UB, resulting in decreased tubule thinning.

### Deleting α-parvin results in excessive F-actin in collecting ducts

The glial cell line-derived neurotrophic factor (GDNF)/Ret receptor signaling axis is required for early UB branching and elongation (40, 41). To address if this signaling was perturbed, we measured Ret mRNA expression using RNA scope and found no difference in Ret expression between Hoxb7:α-pv^f/f^ and α-pv^f/f^ E15.5 kidneys. (Fig. S4). As α-parvin can regulate cell-cell junctions (42), we assessed whether there were polarity defects in the Hoxb7:α-pv^f/f^ CDs by examining ZO-1 (Fig 4A) and E-cadherin (Fig 4B) distribution using immunofluorescence. No abnormalities in localization or amount of these polarity markers were noted. In addition, the localization of the integrin β1 subunit (Fig 4C) and the basement membrane as depicted by laminin staining (Fig 4D) were normal. Thus, the phenotype was not caused by polarity defects. We next investigated the actin cytoskeleton by culturing isolated E11.5 kidneys and whole mount E-cadherin staining was used to identify the UB (Fig 4E a, a’). The actin cytoskeleton was stained with rhodamine-conjugated phalloidin, after which the tissue was subjected to super-resolution confocal microscopy (Fig 4E b-f, b’-f’). Analysis of actin distribution was conducted using a 1 μm z-step across the entire UB tip. We reconstructed the images computationally in 3D to highlight the tubule outline (Fig 4E b, b’). An enlarged tip was evident at the end of a slender trunk in the α-pv^f/f^ UB. However, there was no discernible trunk, and the tip was notably larger in the Hoxb7:α-pv^f/f^ UB (Fig 4E c, c’). Actin filaments were observed on both the apical and basal-lateral borders of the UB cells of the α-pv^f/f^ and Hoxb7:α-pv^f/f^ mice, but they appeared thicker on the basal side of Hoxb7:α-pv^f/f^ CD cells compared to those of α-pv^f/f^ CD cells (Fig. 4E d and d’). When we specifically examined the basal side of the CD cells in super-resolution, we found a clear circumferential actin belt in the α-pv^f/f^ UB cells (Fig. 4E e and f). By contrast, pronounced filamentous actin (F-actin) bundles were observed on the basal side of Hoxb7:α-pv^f/f^ UB tip (Fig 4E e’ and f’). These data suggest that there is excessive F-actin on the basal sides of UB cells, which suggests that α-parvin promotes actin depolymerization in these epithelial cells.

**Figure 4.**
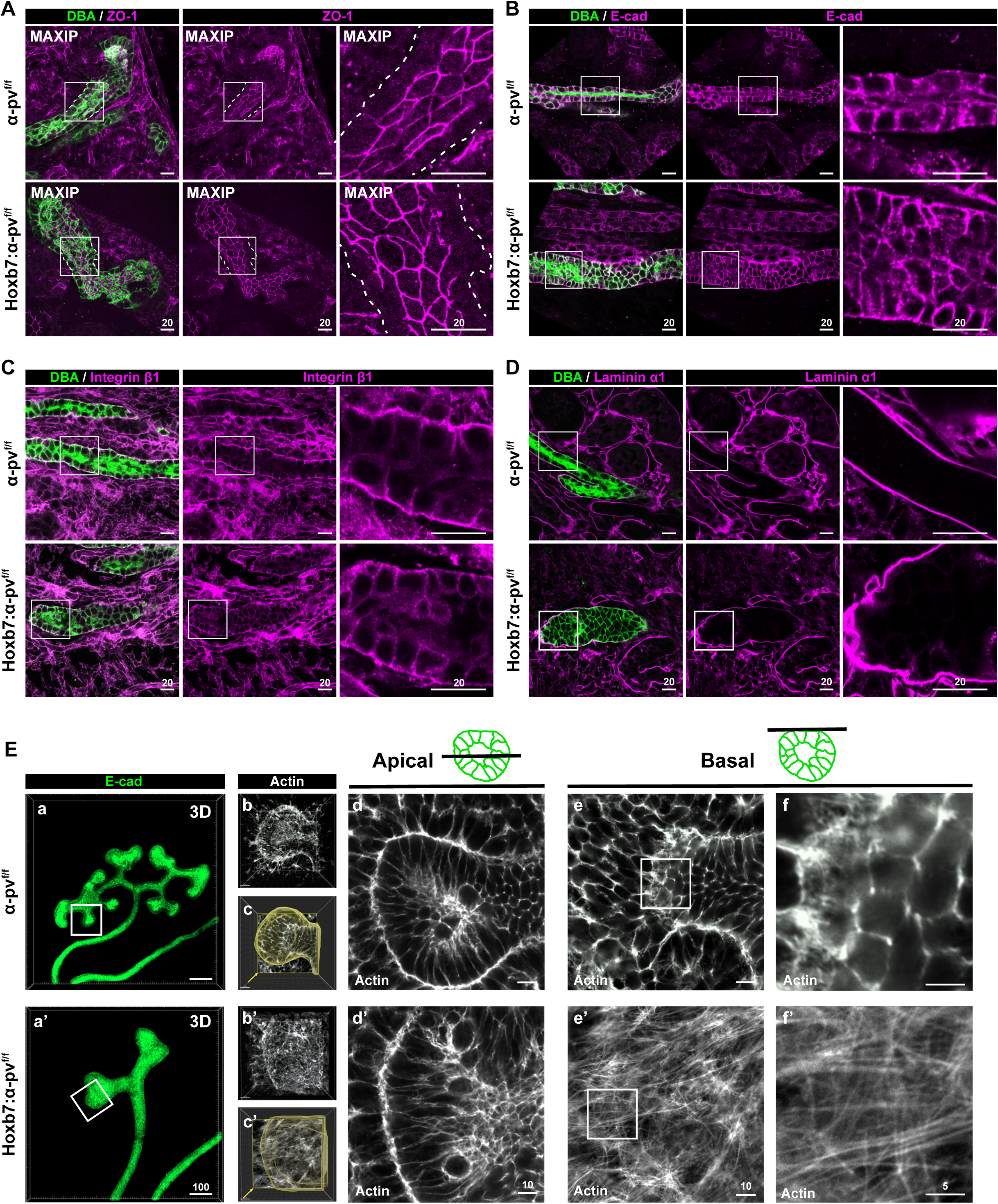
Deleting α-parvin in the UB causes excessive F-actin formation on the basal side of CDs. (A-D) The CDs of E18.5 α-pv^f/f^ and Hoxb7:α-pv^f/f^ mice were labelled with DBA and stained for ZO-1 (A), E-cadherin (B), integrin β1 (C) or laminin α1 (D) and imaged with confocal microscopy. ZO-1 images were shown as maximum intensity projections (MAXIP) to visualize apical junctions. A white dashed line is used to outline the basal side of the tubule. Images are representative of three mice per genotype analyzed. **(E)** E11.5 kidneys isolated from α-pv^f/f^ and Hoxb7:α-pv^f/f^ mice were cultured for 12h and labelled with E-cadherin (green, 3D rendering) (a and a’) and stained with rhodamine-phalloidin (white) (b-f and b’-f’) after which they were subjected to confocal microscopy analysis. 3D reconstruction of one branching tip is shown (b, b’). The tip surface (outlined) was subjected to 3D rendering using Imaris 10.0 (c, c’). Apical (d, d’) and basal (e, e’) actin distribution is shown. Basal F-actin stress fibers are demonstrated at higher magnification (f, f’) in the Hoxb7:α-pv^f/f^ mice. Images are representative of three mice per genotype that were analyzed.

### α-parvin^-/-^ CD cells have a severe in vitro tubulogenesis defect

To investigate the cellular mechanisms whereby α-parvin regulates the actin cytoskeleton, we isolated α-pv^f/f^ CD cells from 8-week-old α-pv^f/f^ mice and deleted α-parvin in vitro using adenovirus-mediated delivery of a Cre recombinase as previously described (43). α-parvin deletion (α-pv^-/-^) was confirmed by immunoblotting (Fig 5A) and immunofluorescence (Fig S5A). As we previously showed that deleting ILK in CD cells results in diminished α-parvin but normal Pinch expression, we determined ILK and Pinch expression in these cells. In contrast to the ILK^-/-^ CD cells (24), there was no difference in levels of either ILK or Pinch in α-pv^f/f^ and α-pv^-/-^ CD cells and they were localized in a focal adhesion (FA)-like pattern at the peripheral end of actin fibers (Fig. S5B and C). We also assessed the effects of deleting α-parvin on the formation of the IPP complex in CD cells. Immunoprecipitation of α-parvin in α-pv^f/f^ CD cells verified that the IPP complex was formed in control cells but as expected, ILK and Pinch were not pulled down in α-pv^-/-^ CD cells (Fig 5B). When we performed ILK immunoprecipitation, Pinch was pulled down in both α-pv^f/f^ and α-pv^-/-^ CD cells and there was more Pinch binding to ILK in the α-pv^-/-^ CD cells (Fig 5C and D).

**Figure 5.**
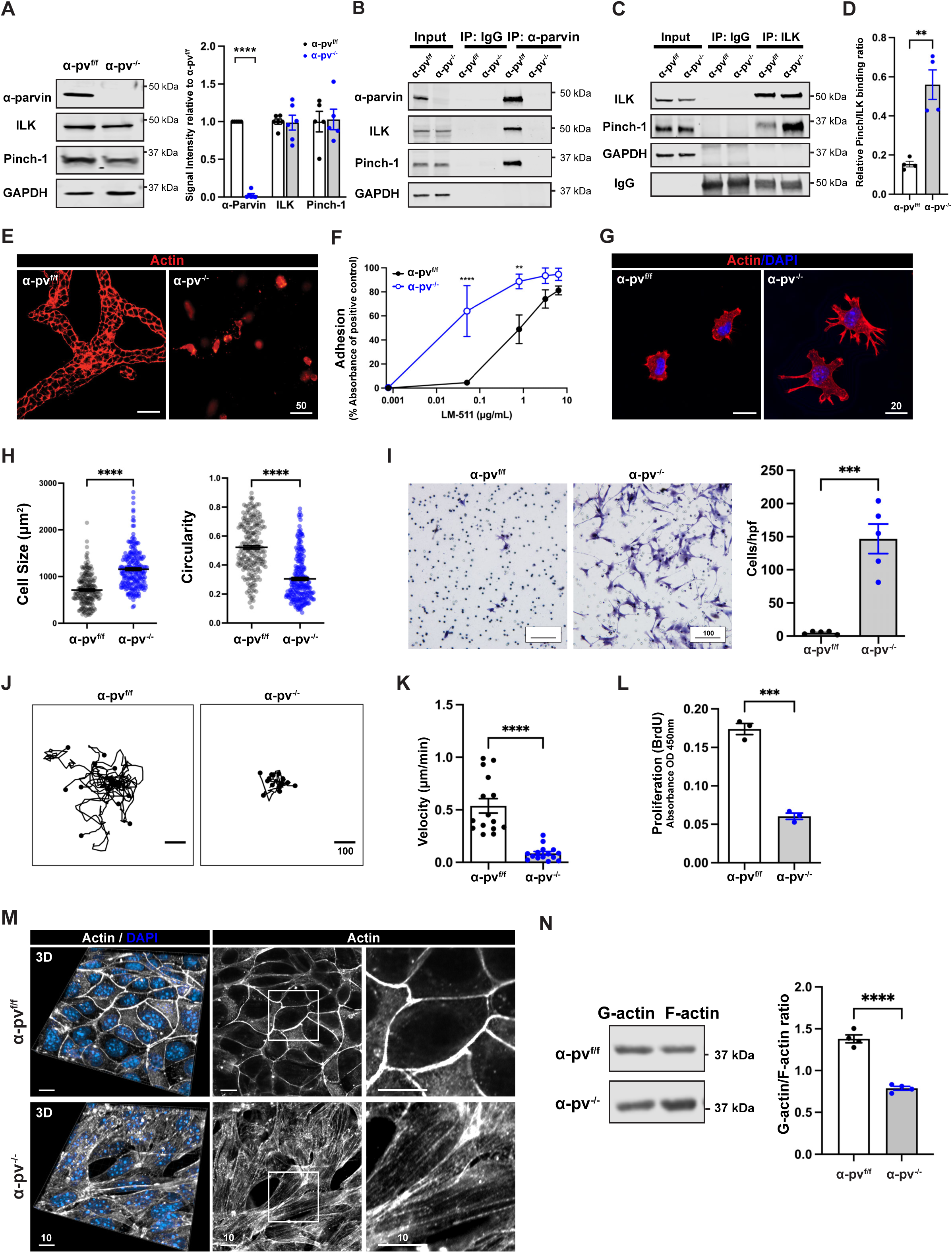
α-pv^-/-^ CD cells have normal ILK-Pinch complex formation but defects in tubulogenesis, adhesion, migration, and spreading due to increased F-actin formation. **(A**) Immunoblots for α-parvin, ILK and Pinch were performed on total cell lysates of α-pv^-/-^ and α-pv^f/f^ CD cells. The relative ratio of α-parvin, ILK and Pinch in the α-pv^-/-^ and α-pv^f/f^ cells was quantified and shown (right panel). Values are mean ± SEM of at least five independent experiments. ****p < 0.0001; 2 tailed t-test. **(B-D)** Cell lysates from α-pv^-/-^ and α-pv^f/f^ CD cells were immunoprecipitated with anti-α-parvin antibody (B) or anti-ILK antibody (C) and the immunoprecipitated proteins blotted for α-parvin, ILK and Pinch. GAPDH was used as a loading control. The relative ratio of Pinch binding to ILK in the α-pv^-/-^ CD cells and α-pv^f/f^ CD cells was quantified and shown graphically (D). Values are mean ± SEM. n = 4, ** p < 0.01; 2 tailed t-test. **(E)** α-pv^-/-^ and α-pv^f/f^ CD cell tubules were grown in 3D collagen-1/Matrigel gels as described in the methods. The tubules were stained with rhodamine-phalloidin and imaged by confocal microscopy. **(F)** α-pv^-/-^ and α-pv^f/f^ CD cell adhesion to different concentrations of laminin-511 at 1 h was measured. Three independent experiments were performed (mean ± SEM), **p < 0.01, ****p < 0.0001; Two-way ANOVA followed by Tukey’s test. **(G)** α-pv^-/-^ and α-pv^f/f^ CD cells were plated on laminin-511 and allowed to spread for 1 h, after which they were stained with rhodamine-phalloidin and DAPI. **(H)** α-pv^-/-^ and α-pv^f/f^ CD cell adhesion area (left) and circularity (right) of (G) was measured using ImageJ. Shown are mean ± SEM of n > 100 cells per group from a single experiment, which is representative of three consistent independent experiments, ****p < 0.0001; 2 tailed t-test. **(I)** α-pv^-/-^ and α-pv^f/f^ CD cells were plated on transwells coated with laminin-511 (10 µg/mL) and allowed to transmigrate through 8 um pores for 4 h. Cells were counted in 5 high-power fields per group. Five independent experiments were performed (mean ± SEM), ***p < 0.001; 2 tailed t-test. **(J-K)** α-pv^-/-^ and α-pv^f/f^ CD cells were plated on slides coated with laminin-511 (10 μg/mL) and cell migration was tracked for 7 hours using ImageJ Manual Track. Individual cell tracks (I) of 15 individual cells per group are shown (I) and used for velocity analysis (J). Shown are mean ± SEM, ****p < 0.0001; 2 tailed t-test. **(L)** α-pv^f/f^ and α-pv^-/-^ CD cells were plated on laminin-511 in the presence of 2% FBS in the presence of BrdU. BrdU incorporation was measured as described in the Methods. Three independent experiments were performed (mean ± SEM), ***p < 0.001; 2 tailed t-test. **(M)** α-pv^f/f^ and α-pv^-/-^ CD cells were grown to confluent monolayers on transwells, stained with rhodamine-phalloidin and analyzed by confocal microscopy. 3D reconstructions (left panels) and cross sections (mid and right panels) are shown. **(N)** G-to-F-actin ratio assay. Representative Western blots for G-actin and F-actin content in α-pv^f/f^ and α-pv^-/-^ CD cells (see Methods). Densitometric analysis shows a significant decrease in the ratio of G-actin relative to F-actin. Four independent experiments were performed (mean ± SEM), ****p < 0.0001; 2 tailed t-test.

Next, we analyzed tubulogenesis in vitro by plating cells into collagen/Matrigel gels. The α-pv^f/f^ CD cells organized into branched tubules after 7 days of culturing, however the α-pv^-/-^ CD cells remained as single cells (Fig 5E). To explain this phenotype, we assessed the ability of α-pv^-/-^ CD cells to adhere, spread, migrate, and proliferate on ECM. Contrary to expectations, α-pv^-/-^ CD cells exhibited a marked increase in adhesion to laminin 511 (Fig 5F) or collagen I (Fig S5D), when compared to α-pv^f/f^ CD cells. They also spread significantly more and formed pronounced filopodia which led to a decrease in circularity when placed on laminin 511 (Fig 5G and H) or collagen I (Fig S5E). They developed significantly larger focal adhesions (FAs), although the number of FAs was not different between the two cell types (Fig S5F and G). Consistent with this data, α-pv^-/-^ CD cells migrated significantly more than α-pv^f/f^ CD cells through transwells coated with laminin 511 (Fig 5H) or collagen I (Fig S5I). However, when the cells were plated on collagen I-coated glass slides and cell movement was tracked, the α-pv^-/-^ CD cells moved less (Fig 5J and K). Also consistent with the inability of the α-pv^-/-^ CD cells to form tubules in 3D-matrices, a severe proliferation defect was evident when these cells were plated on collagen I (Fig 5L).

Due to the abnormalities in cell spreading, adhesion and motility as well as excess basal F-actin formation in the in vivo kidney, we hypothesized that α-pv^-/-^ CD cells would have a defect in the actin cytoskeleton. We therefore stained F-actin in CD cells polarized on transwells with rhodamine phalloidin and visualized them by confocal microscopy. The α-pv^f/f^ cells formed normal actin belts at the cell-cell junction, while α-pv^-/-^ CD cells had excessive F-actin bundles (Fig 5M). When we measured the relative amounts of G versus F actin, there was a decreased G-to-F-actin ratio in the α-pv^-/-^ CD cells (Fig 5N). Taken together, these data indicate that α-parvin enhances epithelial tube formation by regulating actin turnover during cell motility rather than by mediating adhesion to ECM.

### Deleting α-parvin does not affect integrin-dependent signaling

Since integrins are required for cell adhesion, regulate adhesion-dependent signaling pathways and bind to the IPP complex, we initially assessed whether deleting α-parvin altered integrin surface expression by performing flow cytometry of the β1 integrin subunit, and its major binding partners α1, α2 (collagen binding receptors) and α6 (laminin binding receptor). We found no differences between α-pv^f/f^ and α-pv^-/-^ CD cells (Fig S6A). We next performed in vitro replating assays to assess the activation of focal adhesion kinase (FAK), paxillin, Akt, and ERK signaling in α-pv^-/-^ cells. Adhesion of both α-pv^f/f^ and α-pv^-/-^ CD cells induced a significant increase in phosphorylation of these kinases, however there were no major differences between the two genotypes (Fig S6 B, C). Thus, the increased cell adhesion, spreading and migration as well as the decreased random migration observed in the α-pv^-/-^ CD cells are not due to excessive integrin surface expression, activation or integrin-dependent signaling abnormalities and suggest that the defects are primarily within the actin cytoskeleton.

### Deleting α-parvin increases RhoA and Cdc42 and decreases LIMK-dependent cofilin activity

Deleting α-parvin in vascular smooth muscle cells results in actin cytoskeleton defects due to excessive activation of RhoA GTPase (18). We therefore tested the activation status of the Rho family of GTPases (Rho, Cdc42 and Rac1) in α-pv^f/f^ and α-pv^-/-^ CD cells. We found increased activation of Rho A as well as increased expression and activation of Cdc42 (Fig 6A, B) in α-pv^-/-^CD cells. Because the Rho family of GTPases regulates F-actin formation(*44*) by inhibiting the actin depolymerizing factor cofilin (45, 46), we next defined whether α-parvin controls actin dynamics via cofilin. Cofilin phosphorylation at Ser-3 results in inactivation of its actin-binding and depolymerization activity (47). We therefore measured cofilin phosphorylation of α-pv^f/f^ and α-pv^-/-^ CD cells at baseline and after they adhered to collagen I. There was increased cofilin phosphorylation in non-adherent α-pv^-/-^ compared to α-pv^f/f^ CD cells and this difference was accentuated after the cells adhered to collagen I (Fig 6C). Consistent with the data from CD cells, papilla from α-parvin null mice demonstrated increased cofilin phosphorylation (Fig S7A).

**Figure 6.**
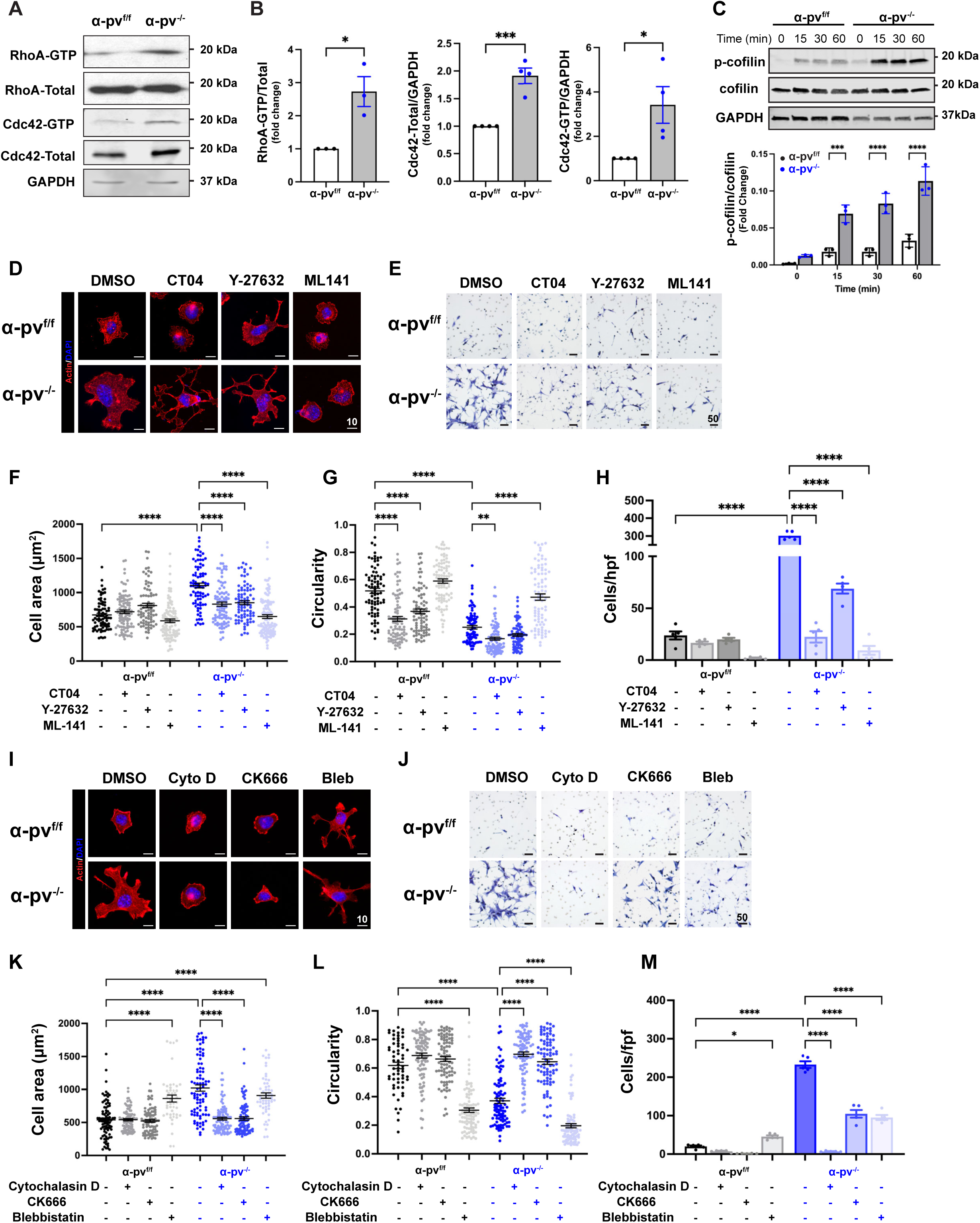
α-pv^-/-^ CD cells have increased RhoA and Cdc42 activity, which regulates cofilin phosphorylation. **(A-B)** α-pv^f/f^ and α-pv^-/-^ CD cells were plated on laminin-511, lysed after 30 min and RhoA or CdC42 activation was evaluated as described in the methods. GTP bound and total RhoA and Cdc42 were analyzed by immunoblotting (C). The ratio of GTP bound and total RhoA and Cdc42 was calculated using densitometry in ImageJ (D-E). The ratio of total Cdc42 in the α-pv^f/f^ and α-pv^-/-^ CD cells was compared (F). GAPDH was used to verify loading. Three-four independent experiments were performed (mean ± SEM), *p < 0.05, ***p < 0.001; 2 tailed t-test. **(C)** α-pv^f/f^ and α-pv^-/-^ CD cells were either left in suspension or plated onto Laminin-511 for 15, 30 and 60 min and immunoblotted for phosphorylated (p-) and total cofilin. GAPDH was blotted to verify equal protein loading. Three independent experiments were quantified using densitometry and shown as individual values and mean ± SEM in the lower panel. ***p < 0.0001, ****p < 0.0001; Two-way ANOVA followed by Tukey’s test. **(D, F and G)** α-pv^f/f^ and α-pv^-/-^ CD cells were treated with the Rho inhibitor CT04 (2 µg/mL), ROCK inhibitor Y-27632 (10 μM) or Cdc42 inhibitor ML141 (10 μM) and allowed to spread on Laminin-511 for 1h. Representative images of rhodamine-phalloidin–stained cells (G) are shown. Spreading area (I) and circularity (J) were quantified using FIJI. Shown are mean ± SEM of n > 100 cells per group from a single experiment, which is representative of three consistent independent experiments, **p < 0.01, ****p < 0.0001; Two-way ANOVA followed by Tukey’s test. **(E and H)** α-pv^f/f^ and α-pv^-/-^ CD cells were treated with Rho inhibitor CT04 (2 µg/mL), ROCK inhibitor Y-27632 (10 μM) or Cdc42 inhibitor ML141 (10 μM) and subjected to a 4h transwell migration assay. Representative images of crystal violet-stained cells (H) are shown. Migrated cells (K) were counted manually in FIJI. Five independent experiments were performed (mean ± SEM) with at least 5 low-power fields per group counted. ****p < 0.0001; Two-way ANOVA followed by Tukey’s test. **(I, K and L)** α-pv^f/f^ and α-pv^-/-^ CD cells were treated with cytochalasin D (100 nM), Arp2/3 inhibitor CK666 (100 nM) or blebbistatin (20 μM) and allowed to spread on laminin-511 for 1h. Representative images of rhodamine-phalloidin–stained cells (A) are shown. Quantification of spreading area (C) and circularity (D) is shown as mean ± SEM of n > 100 cells per group from a single experiment, which is representative of three consistent independent experiments. Five independent experiments were performed and quantified for the migration assays. ****p < 0.0001; Two-way ANOVA followed by Tukey’s test. **(J and M)** α-pv^f/f^ and α-pv^-/-^ CD cells were treated with cytochalasin D (100 nM), Arp2/3 inhibitor CK666 (100 nM) or blebbistatin (20 μM) and subjected to a 4h transwell migration assay. Representative images of crystal violet-stained cells (B) are shown. Migrated cells (E) were counted manually in FIJI. Five independent experiments were performed (mean ± SEM) with at least 5 low-power fields per group counted. *p < 0.05, ****p < 0.0001; Two-way ANOVA followed by Tukey’s test.

We assessed the functional relevance of the increased activity of these Rho GTPases by treating the α-pv^f/f^ and α-pv^-/-^ CD cells with the Rho inhibitor CT04 (C3 exoenzyme analog), the ROCK inhibitor (Y-27632) and the Cdc42 inhibitor (ML141). CT04 and Y-27632 rescued the abnormally increased cell spreading (Fig 6 D, F) and transwell migration of the α-pv^-/-^ CD cells (Fig 6E, H), however the shape (as measured by circularity) of the cells was not rescued (Fig 6G). By contrast, the Cdc42 inhibitor ML141 reduced cell spreading (Fig 6D, F), reverted the abnormal cell shape (Fig 6G) and reduced transwell migration (Fig 6E, H). Cofilin phosphorylation was diminished by decreasing the activity of Rho A (CT104) and ROCK (Y-27632) and this was especially evident when Cdc42 (ML-141) was inhibited (Fig S7 B-D). To examine whether there was cross talk between the Rho and Cdc 42 pathways in mediating their effects on α-pv^-/-^ CD cells, we inhibited both pathways by adding both CT04 and ML141. There was no additive effect on cell spreading, circularity or migration (Fig S7 E-I). As expected, adding both CT04 and ML141 caused severe inhibition of p-cofillin (Fig S7 J). Thus, the Rho GTPases, especially Cdc42, play a key role in α-parvin-dependent regulation of cell shape and movement and this activity is mediated by modulating cofilin-dependent actin-binding and depolymerization.

One of the key mechanisms whereby the Rho family GTPases regulates cofilin activation is via the serine-threonine LIM kinases which directly phosphorylate and inactivate members of the cofilin family (45, 46, 48, 49). To test whether this mechanism accounted for the effects of the increased Rho GTPase signaling found in the α-pv^-/-^ CD cells, we blocked LIM kinase 1 and 2 with the LIMKi 3 inhibitor. This resulted in a decrease in cell area and circularity in both wild type and α-pv^-/-^ CD cells, however the difference was more pronounced in the mutant cells (Fig S8 A-C). A similar result was seen with respect to cell migration (Fig S8 D,E). As expected, the inhibitor markedly degreased cofilin phosphorylation in both α-pv^-/-^ CD cells and wildtype CD cells (Fig S8 F). Taken together these results suggest that the excessive activation of Rho GTPases in α-pv^-/-^ CD cells mediate their effects in a large part by regulating LIM kinase dependent inactivation of cofilin.

### α-parvin-dependent inactivation of cofilin induces excessive actin polymerization

Our data thus far suggests that the cytoskeletal defects observed in the α-pv^-/-^ CD cells are mediated by RhoA and Cdc42 activation, which in turn affects cofilin-dependent actin depolymerization. However, it is possible that the cytoskeletal changes are mediated by abnormal actin-myosin interactions (50, 51). To define which of these mechanisms accounted for the α-pv^-/-^ CD cell phenotype, we examined the effects of cytochalasin D (a cell-permeable potent inhibitor of actin polymerization), CK 666 (an Arp2/3 complex inhibitor) and blebbistatin (a myosin II inhibitor) on CD cell spreading and transwell migration. Cytochalasin D and CK666 decreased the abnormal α-pv^-/-^ CD cell spreading (Fig 6I, K), circularity (Fig 6I, L) and migration (Fig 6K,M) found in α-pv^f/f^ CD cells. By contrast, blebbistatin did not reduce cell spreading (Fig 6I, K) or increase circularity (Fig 6I, L) in α-pv^-/-^ CD cells even though it decreased the transwell migration (Fig 6K,M). Blebistatin did not change cofilin phosphorylation in α-pv^f/f^ or α-pv^-/-^ CD cells (Fig S8G). This data further suggests that the α-pv^-/-^ CD cell abnormalities are primarily due to excessive actin polymerization and not increased myosin contractility.

### α-parvin regulates UB branching morphogenesis by restricting Cdc42 dependent cell re-arrangements and actin polymerization

To assess the functional significance of the excessive activation of the Rho GTPases in UB branching morphogenesis in Hoxb7:α-pv^f/f^ mice in in vivo, we isolated E12.5 UBs from Hoxb7:α-pv^f/f^ and α-pv^f/f^ mice and grew them on transwells for 24 h in the presence or absence of the Cdc42 inhibitor ML141 or the RhoA inhibitor CT104. ML141 partially reversed the branching defect in the Hoxb7:α-pv^f/f^ kidneys (Fig 7A), however there was no rescue of the RhoA inhibitor CT104 (Fig 7B) and there was no additive effect of CT104 when added to ML141 (Fig S9A). This suggests that like in the CD cells, UB branching is facilitated by α-parvin-dependent Cdc42 inhibition.

**Figure 7.**
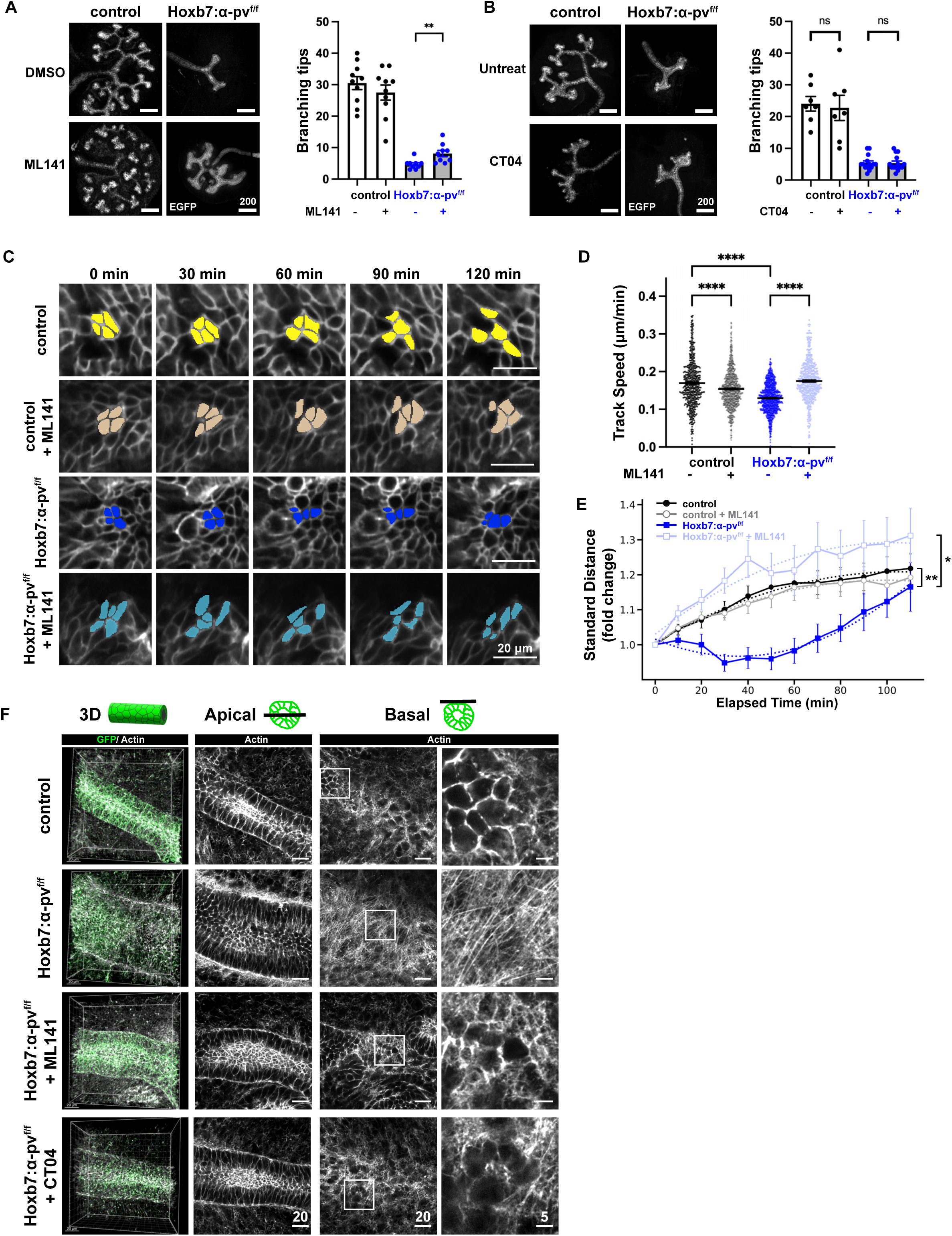
α-Parvin-dependent Cdc42 activity, epithelial cell movement and F-actin turnover is required for UB branching morphogenesis. **(A and B)** E11.5 control (control kidneys were Hoxb7:mTmG^f/f^ unless otherwise noted) and Hoxb7:mTmG^f/f^:α-pv^f/f^ kidneys were isolated and grown ex vivo in the presence of a Cdc42 inhibitor (ML141, 10μM) or a Rho inhibitor (CT04, 2 μg/mL) for 24h and imaged by confocal microscopy. Representative images demonstrate a partial rescue of the branching defect in the Hoxb7:mTmG^f/f^:α-pv^f/f^ kidneys treated with ML141 (A) but CT04 had no effect (B). Branching points were counted in ImageJ with data (mean ± SEM) shown on the right panel. A minimum of 7 embryonic mice were included per group. Each treated kidney was compared with its respective untreated contralateral kidney. The experiments were conducted in three independent sets. **p < 0.001; paired 2 tailed t-test. **(C)** Representative time-lapse images showing the dispersal of a rosette-like cell cluster in control and Hoxb7:mTmG^f/f^:α-pv^f/f^ collecting ducts (CDs) over 2 hours. Cells within the cluster were pseudocolored to visualize cell rearrangement over time. **(D)** Track speed of UB tip cells was significantly reduced in Hoxb7:mTmG^f/f^:α-pv^f/f^ kidneys compared to the control kidneys and was restored by ML141 treatment. Data are shown as mean ± SEM. ****p < 0.0001; 2 tailed t-test. **(E)** Normalized standard distance for the cell clusters were plotted against elapsed time (solid line) with a second-order polynomial curve fit (dotted line). Three independent UB tips per genotype were analyzed, with a minimum of 12 cell clusters from each tip. Data are shown as mean ± SEM. **p<0.01, *p<0.05; Hotelling’s T² tests. **(F)** E11.5 kidneys isolated from control and Hoxb7:mTmG^f/f^:α-pv^f/f^ mice were cultured ex vivo with either the Cdc42 inhibitor ML141 (10 μM) or the Rho inhibitor CT04 (2 μg/mL) for 24 hours and imaged by confocal microscopy. UB trunks were visualized by GFP staining, and actin filaments were visualized with rhodamine-conjugated phalloidin. Panels from left to right show: the 3D reconstruction of the UB trunk, apical section, basal section, and the higher magnification of the basal actin.

To determine the cellular functions regulated by Cdc42 during UB branching morphogenesis, we performed ex vivo cultures of E12.5 kidneys in the presence of the Cdc42 inhibitor ML141, followed by live imaging for 22 hours. Since ML141 significantly reduced membranous GFP intensity in the UB trunk, we limited our analysis to the UB tips, where GFP signal remained robust, likely due to active GFP turnover and synthesis in the proliferating cells (52). Due to the highly motile nature and frequent mitotic events in tip cells, we focused on a shorter, more informative 2-hour window, in contrast to the 8-hour window used for trunk cells (Fig. 3). We found that there was a significant reduction in cell migration speed in Hoxb7:α-pv^f/f^ compared to α-pv^f/f^ tips and ML141 treatment restored cell migration speed to that of α-pv^f/f^ tips (Fig. 7C,D). Interestingly, ML141 modestly decreased migration speed in α-pv^f/f^ kidneys, suggesting that precise regulation of Cdc42 activity is required for normal cell motility in the UB tip. To measure how groups of cells dispersed over time, we followed small clusters of cells in our time-lapse imaging data (see Methods) (Fig. 7C). We found that cell dispersal was significantly reduced in Hoxb7:α-pv^f/f^ compared to α-pv^f/f^ tips (Fig. 7C, E). ML141 treatment restored cell dispersal in Hoxb7:α-pv^f/f^ kidneys back to the levels of α-pv^f/f^ mice. In contrast to its effect on migration speed, ML141 had minimal impact on cell dispersal in α-pv^f/f^ kidneys (Fig. 7C, E), suggesting that cell migration and intercellular movement differ in their sensitivity to low-dose Cdc42 inhibition under normal conditions. Taken together, these results suggest that excess Cdc42 activity contributes to the migration defect in α-parvin–deficient cells, which results in abnormal dispersal dynamics in the α-parvin–deficient UB.

To investigate whether Cdc42 or RhoA activity contributes to the cytoskeletal disorganization observed in Hoxb7:α-pv^f/f^ UBs, we treated isolated E12.5 kidneys with the Cdc42 inhibitor ML141 or the Rho inhibitor CT04 for 24 hours and performed super-resolution confocal imaging (Fig. 7F). The whole explants were stained with phalloidin-647 and the actin was imaged on both the apical and basal side of the entire UB epithelium in 3D. As expected, Hoxb7:α-pv^f/f^ mice exhibited pronounced basal stress fiber formation, with bundled, intersecting F-actin structures visible at the basal side, while control kidneys displayed actin belts and no prominent stress fibers. Strikingly, treatment of α-parvin KO explants with either ML141 or CT04 led to a marked reduction in basal actin stress fiber formation, resulting in an actin architecture that resembled the WT phenotype. Neither inhibitor altered actin organization in control kidneys (Fig. S9B). These results indicate that excessive activation of Rho GTPases, particularly Cdc42 and RhoA, contributes to the aberrant basal actin remodeling observed in the KO, and suggest that α-parvin regulates UB cytoskeletal architecture through modulation of Rho GTPase activity.

## Discussion

In this study we investigated how the scaffold protein α-parvin contributes to UB development, a complex morphological process that is dependent on coordinated cell rearrangement. We utilized ex vivo live imaging to show that α-parvin regulates UB cell reorganization by controlling F-actin depolymerization via the Rho GTPase-dependent LIMK/cofilin pathway. This function contrasts with that of ILK (53), the obligatory binding partner of α-parvin, as well as integrins (54), both of which promote actin polymerization (Fig. 8). These results highlight the importance of adhesion protein-dependent regulation of actin turnover during epithelial cell movement in the setting of branching morphogenesis.

**Fig 8.**
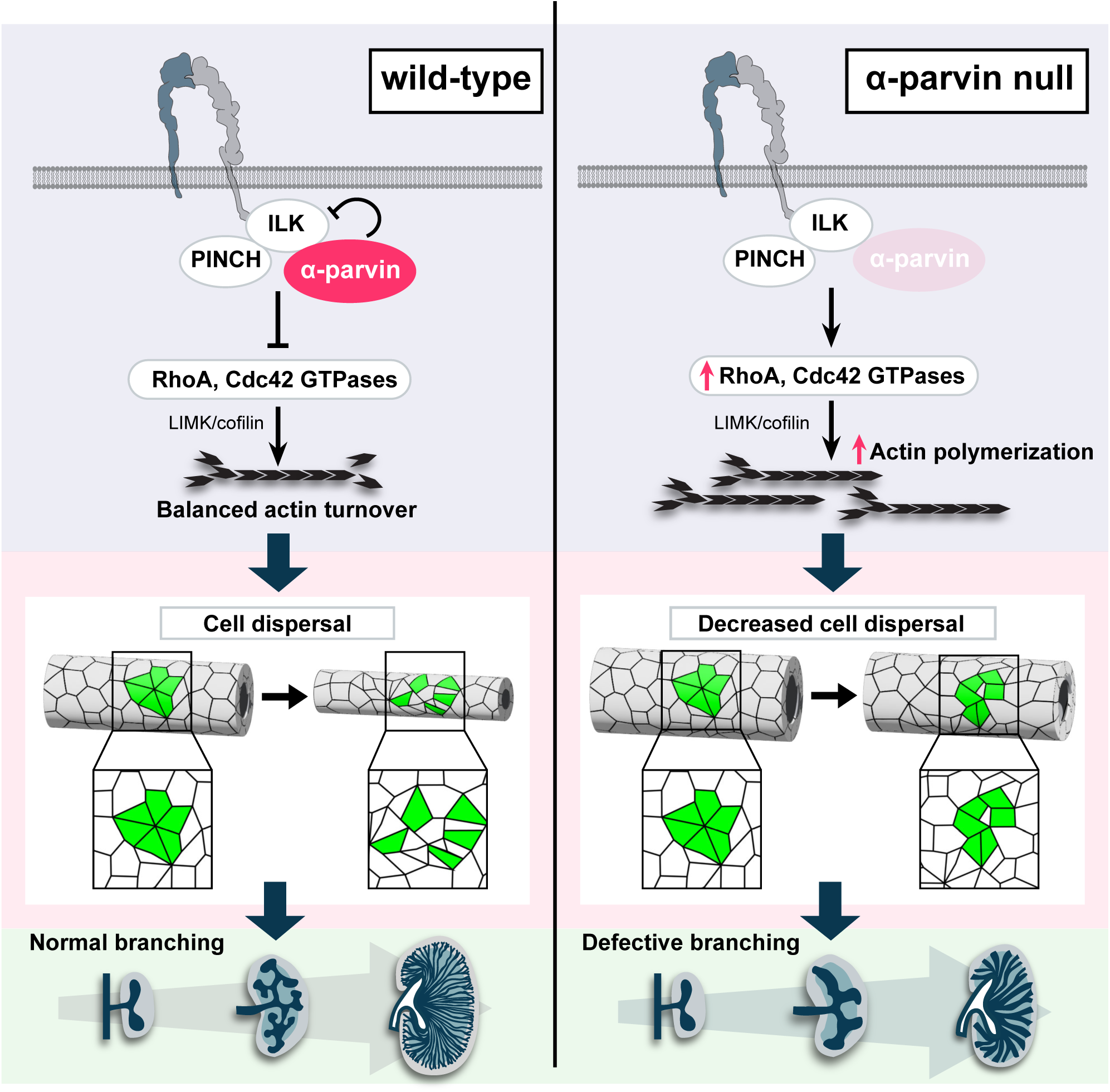

One of the key obstacles to understanding the cellular mechanisms of UB development is the limited live imaging studies performed on this intricate process. By combining an ex vivo kidney explant culture system with a visible membrane tethered GFP and low phototoxicity Airyscan confocal microscopy, we were able to visualize individual cell dynamics in ex vivo branching UB tubules over an extended period. We show that a rosette-based cell intercalation process, occurs in early UB tubule elongation in control mice, like that observed in embryogenesis such as the drosophila germ band epithelium elongation (35) and organogenesis such as Xenopus kidney development (34). We show that growth and thinning of the UB requires cell movement between neighbors and intercalation along an axis that is perpendicular to the direction of elongation, and loss of α-parvin in the developing UB impaired this process leading to less branch formation and decreased tubular narrowing. These defects were due to slower cellular motility and an inability to maintain persistent, coordinated motion across the length of the tissue. This is characteristic of a jammed or solid-like state and is consistent with the slower growth rate of the UB in the α-parvin null mice. Tissue fluidity governs its response to internally generated forces and fluid-like behavior is required for dynamic changes in tissue shape (55, 56). Thus, this change in tissue mechanical properties in combination with the defects in cell morphology and behavior, are sufficient to explain the defects in morphogenesis observed in the α-parvin-deficient UB. The inability of early α-parvin-null UB trunks to narrow was due to actin cytoskeleton abnormalities, rather than a loss of oriented cell division, which is required in later stages of development (9). This finding is consistent with studies conducted in kidneys of mice where both cofilin I and destrin were deleted in the UB using the same Hoxb7 mice (57) and this increased F-actin polymerization together with increased adhesive properties most likely also explains the solidification of the tissue (37). Taken together, our observations suggest that actin-dependent cell movement is a key requirement for early UB morphogenesis.

Since α-parvin is part of the IPP complex that interacts with integrins (19), we expected the α-parvin-UB-null mice, ILK-UB null mice and integrin β1-UB-null mice to have similar phenotypes. While all three mice had a branching morphogenesis defect, there were significant and unexpected differences. The β1 integrin null mice had the most severe branching morphogenesis defect but normal sized tubules (58). The phenotype was due to a severe adhesion defect and a loss of major anchorage-dependent signaling pathways such as Erk and Akt, which are intact in both ILK(24) and α-parvin-null UBs. The ILK-UB null mice have a moderate branching defect and marked tubular obstruction (24); due to CD proliferation within the tubular lumens caused by a loss of p38/MAPK-dependent contact inhibition. The α-parvin-UB- null mice also had a moderate UB branching defect, however the CDs in these mice were broadened due to the inability of epithelial cells to reorganize during UB branching and thinning. These phenotypic differences in the developing UB contrasts with studies in keratinocytes (15, 16), vascular smooth muscle cells (17, 18) and muscle attachment in Drosophila (59, 60) and C.elegans (61, 62) where the mechanisms of action of ILK and α-parvin were shown to be similar. This demonstrates the cell-specific functions of these proteins in different developmental contexts.

It is unclear how the IPP complex is formed and regulates cell function. Deleting ILK results in diminished levels of α-parvin and Pinch and the IPP complex cannot be formed in multiple cell types (13, 19), including CD cells (24, 25). By contrast we demonstrate that deleting α-parvin does not change the levels of ILK and results in increased interactions between ILK and Pinch. This suggests that α-parvin is not an obligatory binding partner of ILK and Pinch, and it might negatively regulate ILK binding to Pinch in CD cells.

One of the key effects of deleting α-parvin in CD cells was an increase in RhoA and Cdc42 GTPase activity. A possible mechanism to explain these data is that the increased Pinch-ILK binding induces Rho GTPase activity through Pinch, as Pinch2 was demonstrated to activate RhoA- and Cdc42 in central nervous system myelination by controlling assembly of the septin protein scaffold (63). It is also conceivable however, that α-parvin reduces Rho GTPase activity in CD cells by recruiting other Rho-regulating scaffold proteins such as cdGAP (64) or ARHGEF6 (23), which have been shown to bind to α-parvin.

In contrast to the increase in RhoA and Cdc42 GTPase activities in α-parvin in CD cells, ILK-null CD cells have decreased RhoA and unchanged Cdc42 and Rac1 activity (Bulus, Zent et al, unpublished data), suggesting that α-parvin and ILK may have antagonistic functions. Consistent with this, the α-Parvin-null CD cells exhibit increased cell spreading, adhesion, and migration, whereas ILK-null CD cells displayed reduced adhesion, spreading and motility (65). Furthermore, p38 phosphorylation is elevated in α-parvin-null cells (data not shown) but diminished in ILK-null cells (65). We attempted to test the genetic interaction between ILK and α-parvin by generating UB–specific double knockouts of ILK and α-parvin, however deleting a single allele of ILK in the α-parvin-null background results in embryonic or early postnatal lethality (Fig S9C). Thus, although we cannot answer whether α-parvin negatively regulates ILK function within the IPP complex, ILK and α-parvin appear to have critical functions that are independent of these interactions.

Another key finding is that the increased adhesion and spreading in the α-parvin CD cells, contrasted with findings in vascular smooth muscle cells, where there was decreased cell adhesion and spreading (18). Although both cell types have increased RhoA activity(18), there was also excessive Cdc42 activation in CD cells. The defect in vascular smooth muscle cells was caused by Rho-dependent elevated MLC phosphorylation resulting in increased actomyosin contractility (18). In α-parvin null CD cells, however, the excess Rho GTPase activation led to decreased actin turnover due to the inhibition of the actin severing function of cofilin. We demonstrate that LIM-kinases partially mediates α-parvin-dependent regulation of cofilin in CD cells. Of note, although the Cdc42 inhibitors completely reconstituted the spreading and migration defects in the CD cells, inhibiting Cdc42 was only partially effective in the organ cultures. This suggests that in addition to the Cdc42-cofillin axis, α-parvin regulates pathways like WASP-mediated actin branching via the Arp2/3 complex (66, 67) and/or formin family proteins(*68*) in in the developing UB.

In conclusion, we demonstrated that cellular rearrangements are critical for UB branching morphogenesis. This process is dependent on dynamic regulation of the actin cytoskeleton by the integrin-binding scaffold protein, α-parvin, which highlights the importance of understanding the complexity of the role of cell-ECM receptors in tubule formation.

## Materials and Methods

### Mice

α-parvin^flox/flox^ mice (α-pv^f/f^) were generated as previously described (18) and were crossed with HoxB7 promoter-driven Cre recombinase transgenic mice (69). All mice are on a C57/B6 background. Female and male mice were used in equal ratios, and aged-matched littermates (α-parvin^flox/flox^; α-pv^f/f^) mice were used as controls. For live imaging, α-pv^f/f^ mice were crossed with mT/mG mice. Mice were bred and maintained under standard conditions in accordance with the Vanderbilt University Institutional Animal Use and Care Committee and conducted in Association for Assessment and Accreditation of Laboratory Animal Care–accredited facilities.

### Antibodies and reagents

#### Antibodies

|  |  |  |
| --- | --- | --- |
| Rabbit anti-α-Parvin (D7F9) | Cell Signaling | 8190 |
| Mouse anti-α-Parvin (clone 1D4) | MilliporeSigma | MABT1441 |
| ZO-1 | Cell Signaling | 8193 |
| Rabbit monoclonal anti-E-cadherin (24E10) | Cell Signaling | 3195 |
| Purified mouse E-cadherin (clone 36) | BD Bioscience | 610182 |
| Rat monoclonal anti-β1-integrin (clone MB1.2) | MilliporeSigma | MAB1997 |
| Rabbit polyclonal anti-SIX2 | Proteintech | 11562-1-AP |
| Laminin | MilliporeSigma | L9393 |
| Rhodamine-conjugated phalloidin | ThermoFisher | R415 |
| DAPI | Cell Signaling | 4083 |
| Ki67 | abcam | ab15580 |
| Rabbit anti-α-Parvin Sepharose Beads | Cell Signaling | 12591 |
| Rabbit monoclonal anti-ILK (4G9) | Cell Signaling | 3856 |
| Mouse monoclonal anti-Pinch | BD Biosciences | 612711 |
| Mouse monoclonal anti-Paxillin | BD Biosciences | 610568 |
| Rabbit monoclonal anti-paxillin (E6R6Z) | Cell Signaling | 50195 |
| Rabbit monoclonal anti-phospho-p44/42 | Cell Signaling | 4370 |
| Rabbit monoclonal anti-p44/42 | Cell Signaling | 4695 |
| Rabbit monoclonal anti-phospho-p38 | Cell Signaling | 4511 |
| Rabbit monoclonal anti-p38 | Cell Signaling | 8690 |
| Rabbit monoclonal anti-phospho-Akt | Cell Signaling | 4060 |
| Rabbit monoclonal anti-Akt (pan) | Cell Signaling | 9272 |
| Mouse monoclonal anti-GAPDH | Cell Signaling | 97166 |
| Rabbit monoclonal anti-β-Actin | Cell Signaling | 4970 |
| Mouse monoclonal anti-β-Actin | Cell Signaling | 3700 |
| Rabbit monoclonal anti-RhoA | Cell Signaling | 2117 |
| Rabbit polyclonal anti-Cdc42 | Cell Signaling | 2462 |
| Rabbit monoclonal anti-phospho-cofilin | Cell Signaling | 3313 |
| Rabbit monoclonal anti-cofilin | Cell Signaling | 5175 |
| Dolichos Biflorus Agglutinin (DBA, FITC-conjugated) | Vector Laboratories | FL-1031 |
| Alexa Fluor 488 goat anti-rabbit | Thermo Fisher | A-11008 |
| Alexa Fluor 488 goat anti-mouse | Thermo Fisher | A-11001 |
| Alexa Fluor 594 goat anti-mouse | Thermo Fisher | A-11005 |
| Alexa Fluor 594 goat anti-rabbit | Thermo Fisher | A-11037 |
| Alexa Fluor 647 goat anti-rabbit | Thermo Fisher | A-21245 |
| Alexa Fluor 647 goat anti-mouse | Thermo Fisher | A-21235 |
| Alexa Fluor 647 goat anti-rat | Thermo Fisher | A-21247 |
| 680RD goat anti-rabbit | LI-COR | 926-68071 |
| 800CW goat anti-rabbit | LI-COR | 926-32211 |
| 680RD goat anti-mouse | LI-COR | 926-68070 |
| 800CW goat anti-mouse | LI-COR | 926-32210 |

#### Chemicals

Prolong gold anti-fade without DAPI (Thermo Fisher; P10144).

M-PER Mammalian Protein Extraction Reagent (Thermo Scientific, 78501)

RIPA (Sigma-Aldrich, R0278)

Rho Inhibitor I (Cytoskeleton, CT04)

ROCK inhibitor (Y-27632, Cell Signaling #13624)

ML141 (Selleck Chemicals, CID-2950007)

Cytochalasin D (Millipore Sigma, C8273)

Arp2/3 inhibitor CK666 (Millipore Sigma, 182515)

Blebbistatin (Millipore Sigma, B0560)

### Kidney Function

Kidney function was determined by measuring serum blood urea nitrogen (BUN) using the QuantiChrom™ Urea Assay Kit (BioAssay Systems, DIUR-100) according to the manufacturer’s protocol. Briefly, serum was mixed with a chromogenic reagent that was measured at 520 nm and the urea concentration deduced from a standard curve.

### Histological analysis

For morphological analysis, kidneys were removed at different stages of development and at the indicated ages in adults. They were fixed in 10% formaldehyde or paraffin and embedded. Paraffin tissue sections (5 µm) were stained with hematoxylin and eosin (H&E) for morphological evaluation by light microscopy. In all experiments, noon of the day on which the mating plug was observed was designated E0.5.

Kidney area was measured in FIJI using cross-sectional sections where the papilla was visible. Kidney length was determined by measuring the longest linear axis across the kidney section. For immunofluorescence, frozen kidney sections were prepared as previously described (70). In brief, kidneys were fixed in 4% PFA for 1hr at room temperature and incubated in 30% sucrose overnight after which they were frozen in OCT for optimal cutting. Sections were blocked with 8% normal horse serum, 0.3% Triton-X100 and 0.1% Tween in PBS for 1 hr and subjected to primary antibodies diluted in the same blocking buffer overnight at 4℃. Samples were washed, incubated with DBA and secondary antibodies, and mounted. Sections were imaged by confocal microscopy (Zeiss LSM 980, Airyscan 2).

For whole mounts, samples were prepared as previously described (41, 71–73). Briefly, embryonic day 10.5-12.5 kidneys were collected, fixed in 4% PFA at room temperature for 1 hr, washed in PBS and blocked in blocking buffer (PBS-BB: PBS with 1% BSA, 0.2% nonfat dry milk powder and 0.3% Triton X-100) (72). Kidneys were incubated with primary antibodies diluted in PBS-BB for 1-3 days (depending on the embryonic stage of the kidneys) at 4℃ and washed overnight with PBST (0.1% Tween) at 4℃. Kidneys were then exposed to secondary antibodies diluted in PBS-BB for 1-3 days at 4℃. Kidneys were washed as described above and examined by confocal microscopy (Zeiss LSM 980, Airyscan 2). Sections were scanned at 1μm steps and Z-stacks were used to conduct 3D reconstruction using FIJI or Imaris. Branching tips were quantified using FIJI software with the following standardized pipeline: images acquired from confocal microscopy were first processed by maximum intensity projection to create a 2D representation. After background subtraction and Gaussian blur filtering to smooth the images, a threshold was applied to generate a binary mask of kidney structures. This mask was then skeletonized, producing a simplified structural representation used to objectively identify branching points and tips. Each terminal endpoint identified in the skeletonized image was counted as one tip.

For tubular diameter analysis, 8-μm-thick frozen sections were used. CDs were labelled with DBA and the entire section was imaged using confocal microscopy (Zeiss LSM980, AiryScan 2). Only diameters of the tubules, where the lumen was visible, were measured. Medians were calculated and significance was determined by the Mann-Whitney U test.

For cell height and number measurement, 40-μm-thick frozen sections were used to obtain Z-stack images of the DBA-labelled collecting tubules. Cortical and medullary CDs were analyzed. The cell number was manually counted in cross sections that were vertically aligned with the tubule’s longitudinal axis, using DAPI as the marker for the nuclei. Cell height was measured using FIJI.

### RNAscope analyses

RNAscope® 2.5 HD Detection Kit-BROWN (Advanced Cell Diagnostics, Cat No: 322330) was used to determine Ret expression pattern in E15.5 kidney. Briefly, E15.5 embryos were fixed in 10% formaldehyde for 24hr at room temperature, paraffin embedded, cut at 5μm, and transferred onto superfrost glass slides. Sample preparation, pretreatment and RNAscope assays were performed according to Formalin-Fixed Paraffin-Embedded (FFPE) Sample Preparation Pretreatment Guide User Manual, Part 1 (Cat No. 322452-USM) and RNAscope® 2.5 HD Detection Kit (BROWN) User Manual, Part 2 (Cat No. 322310-USM). Probe for mouse Ret (Cat No.431791) and negative probe (Cat No. 310043) were used as controls.

### Live-organ Imaging of kidney ex vivo

HoxB7^Cre^ (Hoxb7) or Hoxb7:α-pv^f/f^ were crossed with mT/mG mice (Jackson laboratory, #007676). Kidney explants from E12-12.5 HoxB7^Cre^:mT/mG (Hoxb7), Parvin^fl/wt^:mT/mG:HoxB7^Cre^ (Hoxb7:α-pv^f/wt^) and Parvin^fl/fl^:mT/mG:HoxB7^Cre^ (Hoxb7:α-pv^f/f^) were isolated as mentioned above (71, 72). Ex vivo tissue culture set-up for live imaging was adapted from the FiZD method^64^ and salivary gland live imaging^63^. Briefly, double-adhesive imaging spacers (Grace Bio-Labs, 654008) were sterilized by soaking in 70% ethanol for 3 min and attached to the glass bottoms of 35mm ibidi dishes (ibidi, 81218-200). Kidneys were carefully transferred to the center of ibidi dish using a 200μL pipette. A maximum of 3 kidneys were included in one dish. 10μL of Matrigel (Corning, 356230, 5μg/μL) was dropped onto each kidney to prevent tissues from moving. The transwell filter of the Snapwell (Corning, #3801, 0.4μm pore size) was stuck to the image spacer so that the kidneys were sandwiched between the glass bottom and the filter. 500 μL of organ culture medium (DMEM/F12, 10% FBS, with penicillin/amoxicillin) was then added to the well. The kidneys were incubated at 37 ℃ with 5% CO_2_ for 16 hours before imaging. For inhibitor experiments, kidneys were submerged in inhibitors for 1 hr at 37 ℃ with 5% CO_2_ before being transferred to the ibidi dish. A Zeiss LD LCI Plan-Apochromat 25x, 0.8 NA, water-immersion objective on a Zeiss LSM980 AiryScan 2 Confocal Microscope controlled by Zeiss Zen Blue software was used for image acquisition. Images were acquired at 1μm z intervals at ∼50 μm thickness and 10 min intervals for 22 hrs. using CO-8Y mode. Laser was used at 5.5% for excitation of membrane-EGFP.

### Live video analysis

3D surface reconstruction of indicated time points was performed by drawing polylines along the epithelial surface at every other z-plane (2μm interval) on a x-y view using Imaris software.

To study the change in tubular size during elongation, z-stack images were rotated so that the longitudinal axis of the tubule was aligned with the x-axis in FIJI. The size of the tubule was assessed by measuring the cross-sectional area in the orthogonal view, accomplished by manually drawing polylines around the tubule at indicated time points during an 11-hour time span.

To measure the orientation of cell division, UB trunk cells (50∼100 μm from the tip edge) were manually identified when they were in either telophase or cytokinesis, as observed in the time-lapsed movies. The division angle was measured in FIJI using the “angle” tool as previously described(*74*). Briefly, a first line was drawn connecting the centroids of the dividing cells and the angle was defined by drawing a second line parallel to the longitudinal axis of the tubular lumen. The measurement of cell re-arrangement was adapted from Packard et al(*75*). Briefly, cells were first segmented using Cellpose (cyto3 model) (76), followed by manual correction. Cell trajectories were then tracked using TrackMate (77) in FIJI. For cell dispersal analysis, 4 to 8 cells that shared the same convergent point (defined as located within 15 pixels of one another) were randomly picked and identified as a cell cluster. Those cell clusters were then followed over indicated time. Dispersal was quantified as the standard distance, or the geometric mean of the distance of each cell from the group centroid over time. Standard distance (SD) was calculated using the following function: 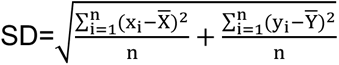, where (x, y) is the pixel coordinate and 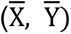 is the mean center of the cells.

To evaluate whether the trend of cell dispersal differed between experimental groups, we analyzed UB trunk and UB tip regions separately due to their distinct dispersal dynamics. For UB trunks, the standard distance over time was fit into a linear regression equation y = at + b, where t is time. The slope (a) and intercept (b) were extracted as descriptors of dispersal behavior. To statistically compare dispersal trends between groups, we performed analysis of covariance (ANCOVA) on the regression slopes. Dispersal behavior at the UB tips followed a nonlinear pattern and was better captured using a second-order polynomial fit. For each cell cluster, the standard distance over time was fitted to the equation y = at² + bt + c. The resulting coefficients (a, b, c) were treated as multivariate representations of dispersal dynamics. To compare these across experimental groups, we performed pairwise Hotelling’s T² tests, a multivariate extension of the t-test. P-values were adjusted for multiple comparisons using the false discovery rate (FDR) method.

### Method for coarse grained velocity field estimation

Coarse grained Particle Image Velocimetry was performed using quickPIV – an opensource script written in Julia Language. The time series of the 3D image stacks were processed in Jupyter notebooks with a window size of (32,32,7) pixels in the (x,y,z) directions respectively. Based on a constant height of the image stacks, the z dimension of the window was set to 7 pixels. Window overlaps of (8,8,3) pixels were specified unless explicitly stated with a search margin set to (10,10,3) pixels to account for the boundaries. To compute the motion similarity maps, a scalar product between the raw velocity vector and its 4-neighbours were used. The visualizations of the velocity vectors and the motion similarity maps were rendered in Paraview 5.10.

### Cell culture

Primary CD cells were isolated from kidneys of α-pv^f/f^ mice and immortalized with SV40 as previously described(*58*). The PARVA gene was deleted by infecting with adenoCre virus, after which null-CD cells were cloned. CD cells were cultured in DMEM/F12 supplemented with 10% FCS and antibiotics. Deletion of α-parvin was verified by Western blot analysis. Numerous clones were tested, and they behaved similarly. For cell treatments, all drugs were dissolved for stock solutions and used according to the manufacturer. Solvents for stock solutions were used as negative control treatments in respective experiments.

### Immunoprecipitation

For immunoprecipitation, cells were lysed with M-PER and protein extracts (0.5 mg for α-Parvin immunoprecipitation and immunoblotting for α-Parvin, Pinch and ILK, and 1.0 mg for ILK immunoprecipitation and immunoblotting for ILK and Pinch) were pre-conjugated beads or pre-cleared with protein A/G beads and incubated with the specific primary antibody or overnight at 4°C with rotating, respectively. The antibody-protein complex was washed twice with TBS-0.1%Tween and twice with TBS-1% TritonX100 and collected by centrifugation. The immune complex was eluted by SDS-PAGE sample buffer and subject to Western blot analysis.

### Tubulogenesis assay

Tubulogenesis of CD cells was performed in 3D ECM gels composed of 1.5mg/mL rat tail collagen I (Corning, 354236) and 0.25mg/mL Matrigel in DMEM containing 20 mM HEPES. CD cells (1.5 x 10^3^) were mixed with the Collagen/Matrigel mixture and transferred into a 96-well plate and returned to tissue incubator (37℃, 5% CO_2_). After the cell/Collagen/Matrigel mixture solidified, 100 μL of DMEM/F12 with 10% FBS and antibiotics were added to each well to prevent evaporation. The cells were allowed to grow for 7 days. The gels were stained with rhodamine-conjugated phalloidin, and the tubules were imaged using a Zeiss Axio 510 confocal microscope (400x).

### Cell spreading and focal adhesion analysis

8-well chamber slides coated with collagen I (10μg/mL) or laminin 511 (10μg/mL) were blocked with 1% heat-denatured BSA for 1 hr. CD cells were then plated in serum-free medium with indicated treatments onto the slides. Cells were allowed to spread for 1 hr and fixed with 4% PFA and stained with rhodamine-conjugated phalloidin. Images were collected with fluorescence microscope (Olympus, 20X/1.25NA) and the cell area and circularity were determined using FIJI software. For focal adhesion (FA) analysis, cells were allowed to spread for 4 hrs. FAs were labelled by immunostaining of paxillin, and the number/cell area and FA size/cell area were quantified using FIJI.

### Cell adhesion, migration, proliferation

For the cell adhesion assay, 96-well round-bottom plates (Corning, #38018) were coated with different concentrations of Collagen I or Laminin 511 and blocked with 1% heat-denatured BSA. 10^5^ cells in serum-free DMEM/F12 were placed in each well for 1 hr; nonadherent cells were removed and the remaining cells were fixed with 4% PFA, stained with 1% crystal violet, and solubilized in 20% glacial acid. The plates were measured for absorbance at 540nm.

For the transwell migration assay, transwells with 8-μm pore size (Corning, #3422) were coated with Collagen I or Laminin 511 (10 μg/mL) on the lower side of the filter, and 10^5^ cells were added to the upper well in serum-free medium. For inhibitor treatments, cells were pre-treated with Y-27632 (ROCK inhibitor) and ML141 (Cdc42 inhibitor) for 45 minutes before seeding. CT04 (Rho inhibitor), LIMKi 3 (LIMK inhibitor), blebbistatin, cytochalasin D, and CK666 were mixed with the cell suspension at seeding. Cells that migrated through the filter after 4 hrs. were stained with 1% crystal violet (Sigma-Aldrich, C3886) and hematoxylin Gill #3 (Sigma-Aldrich, GHS332) for cell nuclei. Images were collected with microscope (Olympus, 20X/1.25NA) and 5 random fields were analyzed per group. Cells were counted using FIJI.

For the random migration assay, cells were sparsely seeded on Collagen I (10 μg/mL)-coated 35 mm glass-bottom ibidi dishes (ibidi, 81218-200) in serum-free culture medium. Time-lapse DIC imaging was conducted at 10-minute intervals for 24 hours using a 10x objective lens on a Zeiss LSM980 confocal microscope. DIC images were acquired with a laser power of 7%, using a PMT gain of 700V. Cell tracking was performed using TrackMate(*77*) in FIJI, measuring cell velocity as mentioned above.

Cell proliferation was assessed by measuring incorporation of 5-bromodeoxyuridin (BrdU) in an enzyme-linked immunosorbent assay according to the manufacturer’s manual (Exalpha, X1327K). Briefly, 96-well plates were either left uncoated, or coated with Collagen I or Laminin 511 (50 μg/mL) and blocked with 1% BSA. 1500 cells were plated in medium (DMEM/F12 with 2% FBS) each well and allowed to adhere for 6 hrs. and then incubated with BrdU for 16 hrs. BrdU incorporation was quantified by a change of absorbance (OD) at 450 nm.

### Integrin-related signaling pathway and cofilin activation analysis

For adhesion-dependent integrin signaling and the cofilin activation assay, serum-starved CD cells were plated in serum-free medium on Collagen I (10 μg/mL) for the indicated time points. For inhibitor treatments, cells were pre-treated with Y-27632 (ROCK inhibitor) and ML141 (Cdc42 inhibitor) for 45 minutes before seeding. CT04 (Rho inhibitor), LIMKi 3 (LIMK inhibitor), blebbistatin, cytochalasin D, and CK666 were mixed with the cell suspension at seeding. Non-adherent cells were removed by gently washing with PBS. Adherent cells were lysed using M-PER reagent with protease and phosphatase inhibitor cocktails. Protein extracts (20-40 μg) were subject to western blot analysis.

### Flow cytometry

Flow cytometry was performed as previously described(*78*). CD cells were incubated with the indicated antibodies followed by fluorescein isothiocyanate-conjugated secondary antibodies.

### Rho GTPases Pull down and activity assays

The activation status of RhoA, Cdc42, and Rac1 was assessed using pull-down assays specific for the GTP-bound form of the Rho GTPases. We used a Rho Activation Assay kit (Millipore Sigma, 17-294) according to manufacturer’s protocol. The pull downs were quantified by Western Blots.

### G-to-F-actin ratio assay

G-actin/F-actin ratio analysis was performed using the G-actin/F-actin in vivo assay Beachem kit (Cytoskeleton, BK037). Briefly, equal amount of protein was determined by BCA protein quantification prior to the G-to-F-actin ratio assay. Cells were lysed in F-actin stabilization buffer containing 1 mM ATP, followed by centrifugation to separate the F-actin (pellet) and G-actin (supernatant) pools. The F-actin pellet was then dissolved in F-actin depolymerization buffer which contains 8M urea. The resulting fractions were subject to SDS-PAGE and actin was quantified by Western blot.

### In vitro Immunofluorescence

For cellular immunostaining, cells were grown on 8-well Milli cell EZ chamber slides (Millipore, PEZZGS0816) coated with 10 µg/ml collagen I or 10 µg/ml Laminin 511 and allowed to adhere and spread for the times indicated. Cells were fixed in 4% PFA, rinsed with PBS and permeabilized and blocked with blocking buffer (0.3% Triton X-100, 8% normal horse serum in TBS. Subsequently, cells were incubated with the indicated primary antibodies followed by secondary antibodies. The coverslips were then washed and mounted on glass slides using prolong gold anti-fade mounting media. Fluorescent images were acquired using a Zeiss laser scanning confocal microscope (LSM980, 63X/1.5NA oil Olan Apochromat oil). For polarized cell staining, cells were grown on transwell inserts until they formed a monolayer, fixed in 4% PFA, permeabilized, and blocked as mentioned above. The cells were then stained with Rhodamine-conjugated phalloidin. Z-stack images were acquired at 1μm intervals. Line scan was performed using the plot profile tool of FIJI software.

### Ex vivo kidney cultures with inhibitors

Embryonic 12.5-day kidney explants with mT/mG transgene were collected as described above. Kidneys were submerged in inhibitors 1 hr before transferring to a Transwell (Corning, #3460). Kidneys were cultured with complete medium (DMEM/F12, 10% FBS, antibiotics) with inhibitors for 24 hrs. and fixed. Images were examined by confocal microscopy (Zeiss LSM 980, Airyscan 2). Sections were scanned at 1μm steps and maximal intensity projection was used for visualization and analysis in FIJI. Branching was assessed by quantifying the number of tips from images.

### Statistics

Data are shown as mean ± SEM. Unpaired two-tailed t test was used to evaluate statistically significant differences (*, p < 0.05) between two groups. Two-way ANOVA was used to test statistical significance (*, p < 0.05) among multiple groups. Post-hoc comparisons of ANOVA were corrected with the method of Tukey. For fold-change calculations (e.g., Western blot densitometry), the control number was normalized to 1 for each comparison. Data distribution was assumed to be normal, but this was not formally tested. GraphPad Prism software was used for statistical analysis.

## Acknowledgements

We are grateful to the cell imaging shared resource (CISR), our flow cytometry cores at Vanderbilt and the Nashville VA, and our pathology core (Translational Pathology Shared Resource), and we would like to thank J. Schafer, T. Vadakkan C. Warren, B. Matlock, and D. Flaherty for technical assistance.

## Funding

National Institutes of Health grant R01 DK069921 (R.Z.)

National Institutes of Health grant R01 DK088327 (R.Z.)

National Institutes of Health grant R01 DK127589 (R.Z.)

National Institutes of Health grant K08 DK134879 (F.B.)

National Institutes of Health grant R01 DK119212 (A.P.)

Veteran Affairs Merit award I01-BX002196 (R.Z.)

Veteran Affairs Merit award 1I01BX002025 (A.P.)

American Society of Nephrology Kidney Cure Pre-Doctoral Fellowship (X.D.)

American Society of Nephrology Kidney Cure Ben J. Lipps Fellowship (F.B.)

Keck Foundation grant (R.Z.)

Max Planck Society (S.W.)

Academy of Finland Center of Excellence BarrierForce grant # 346131 (S.W.)

National Institutes of Health grant P30-CA068485 (Vanderbilt Cell Imaging Shared Resource) Flow cytometry experiments were performed in the Nashville VA flow cytometry core (U.S. Department of Veterans Affairs, Tennessee Valley Healthcare System, Nashville, TN) and the VMC Flow Cytometry Shared Resource (supported by the Vanderbilt Ingram Cancer Center, P30-CA68485, and the Vanderbilt Digestive Disease Research Center, DK058404).

National Cancer Institute/National Institutes of Health Cancer Center support grant P30-CA068485 (Translational Pathology Shared Resource

## Contributions

Conceptualization: XD, FB, NB, AP, RZ

Methodology: XD, FB, AH, NB, GM, GB, WZ, OV

Investigation: XD, AH, NB, GM, GB, SL, WZ, MM, KB, CM, OL

Visualization: XD, AH, SW

Supervision: RZ, SW

Writing-original draft: XD, SW, RZ

Writing: review & editing: XD, FB, AP, SW, RZ

## Data and materials availability

All data are available in the main text or the supplementary materials

