## Supplementary figures1-9 for "α-Parvin regulation of cell re-arrangement is critical for ureteric bud branching morphogenesis"

Fig.S1

A

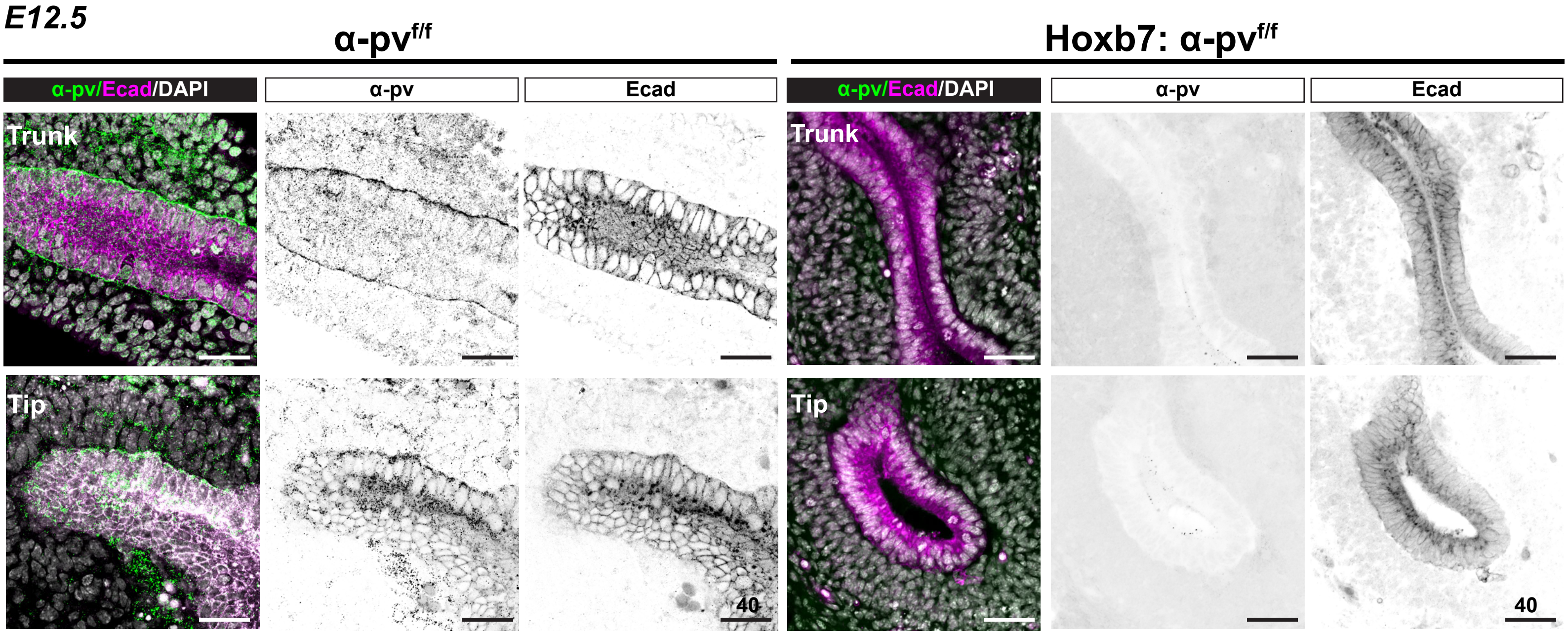


B

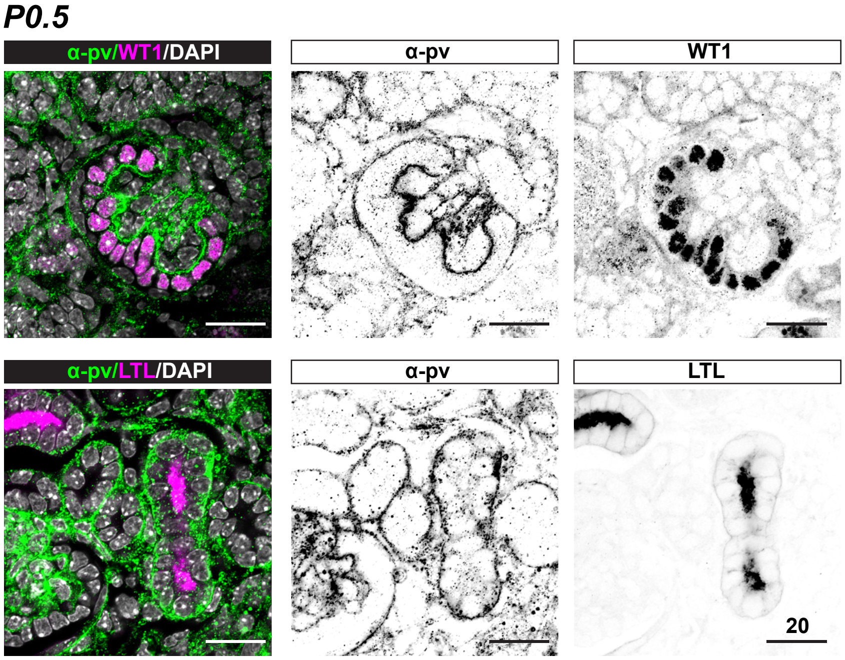


C

*$\alpha$-pv$^{f/f}$ X Hoxb7:$\alpha$-pv$^{f/+}$*

| | $\alpha$-pv$^{f/f}$ | Hoxb7: $\alpha$-pv$^{f/f}$ | $\alpha$-pv$^{f/+}$ | Hoxb7: $\alpha$-pv$^{f/f}$ |
|---|---|---|---|---|
| **Expected** | 25% | 25% | 25% | 25% |
| **Observed embryos** | 22.5% (n=18) | 25% (n=20) | 21.25% (n=17) | 21.25% (n=25) |
| **Observed pups** | 27.7% (n=41) | 29.1% (n=43) | 19.6% (n=29) | 23.6% (n=35) |

D

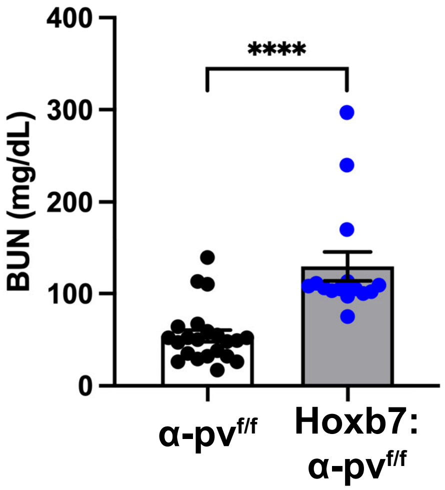

Fig S2

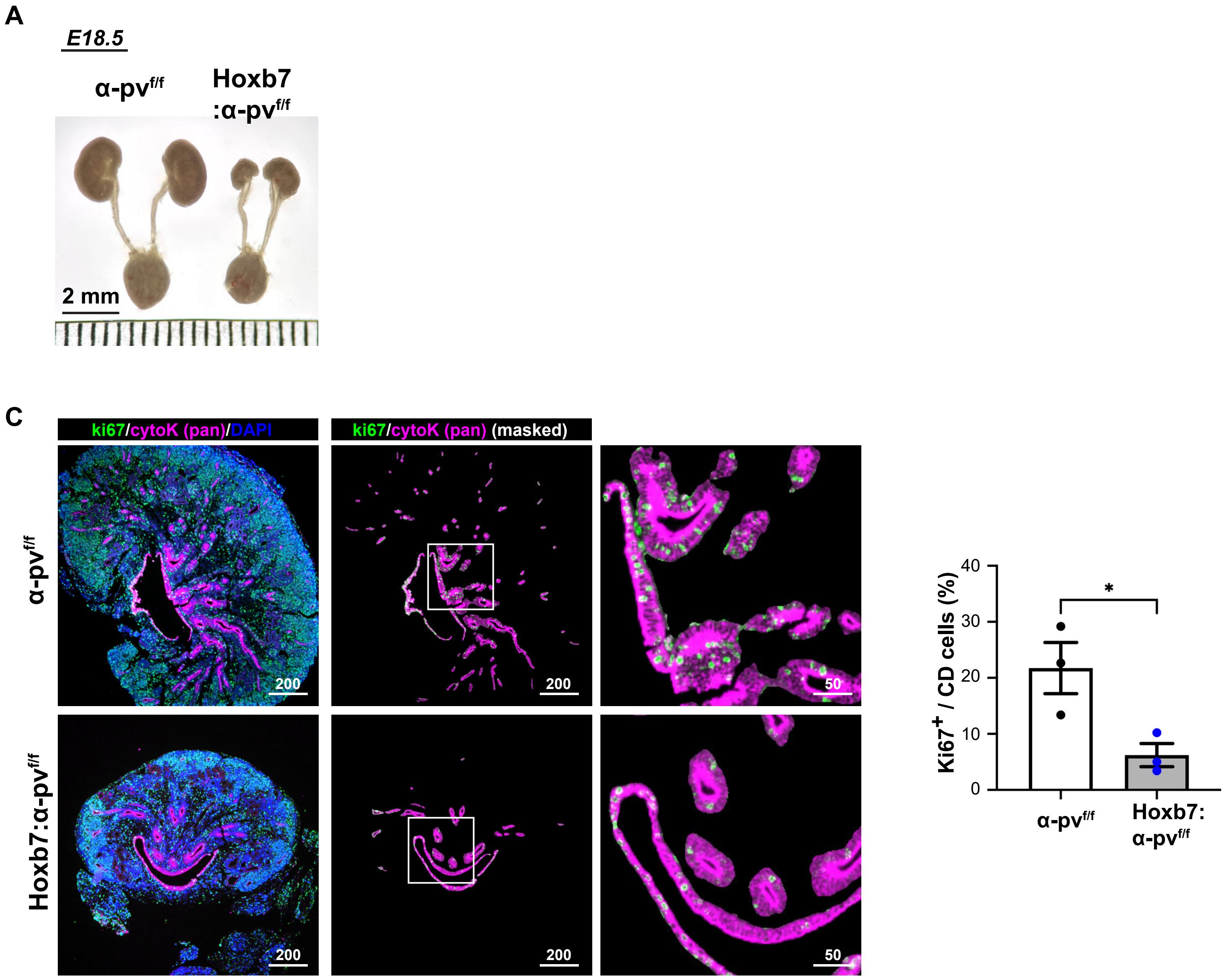
A
E18.5
α-pvf/f
Hoxb7
:α-pvf/f
2 mm
C
ki67/cytoK (pan)/DAPI
ki67/cytoK (pan) (masked)
α-pvf/f
Hoxb7:α-pvf/f
200
50
Ki67+ / CD cells (%)
40
30
20
10
0
*
α-pvf/f
Hoxb7:
α-pvf/f

Fig S3

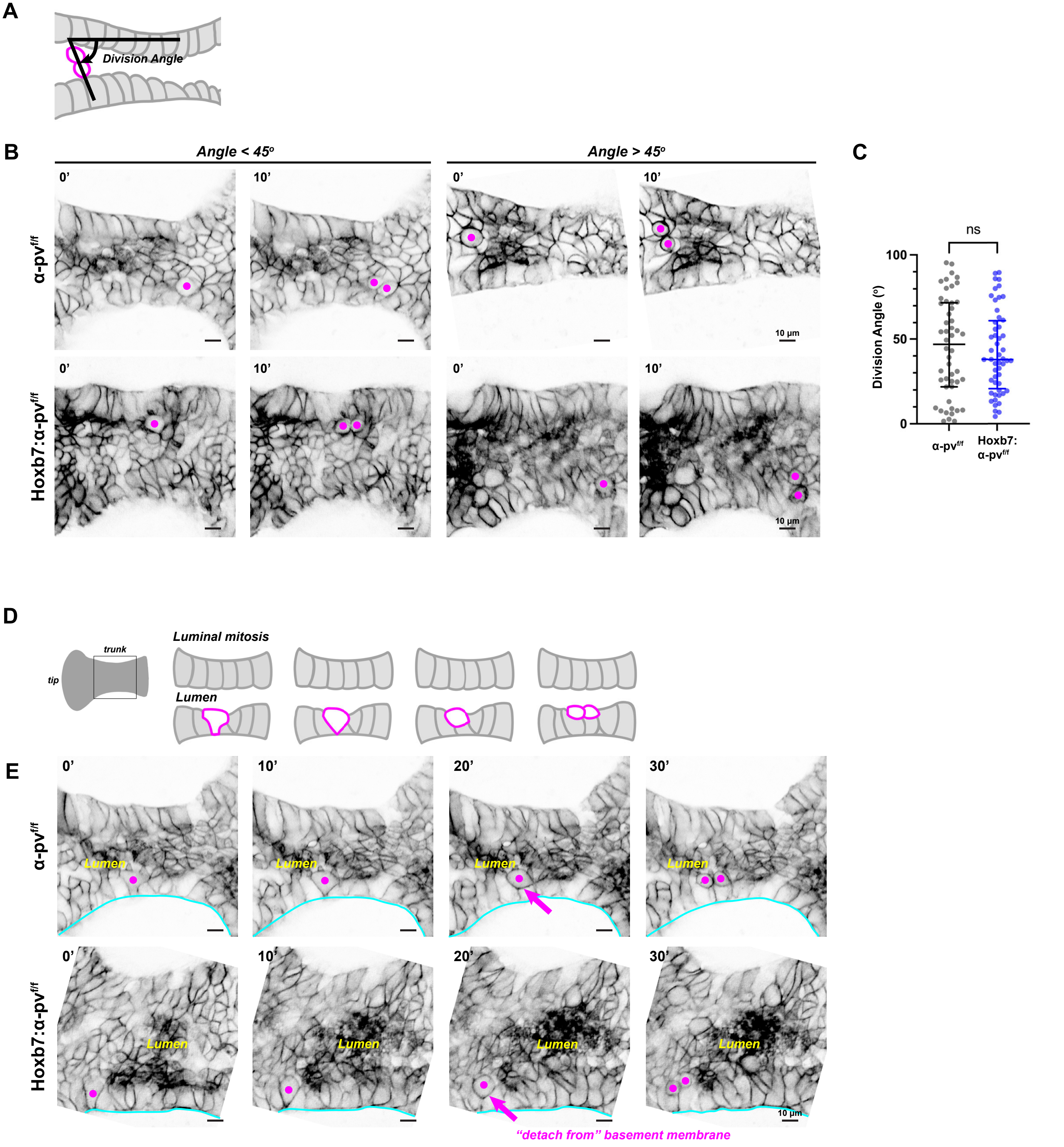
A
Division Angle
B
Angle < 45º
Angle > 45º
0'
10'
α-pv^f/f
Hoxb7:α-pv^f/f
10 µm
C
ns
Division Angle (º)
100
50
0
α-pv^f/f
Hoxb7:
α-pv^f/f
D
trunk
tip
Luminal mitosis
Lumen
E
0'
10'
20'
30'
Lumen
"detach from" basement membrane

Fig S4

**A**

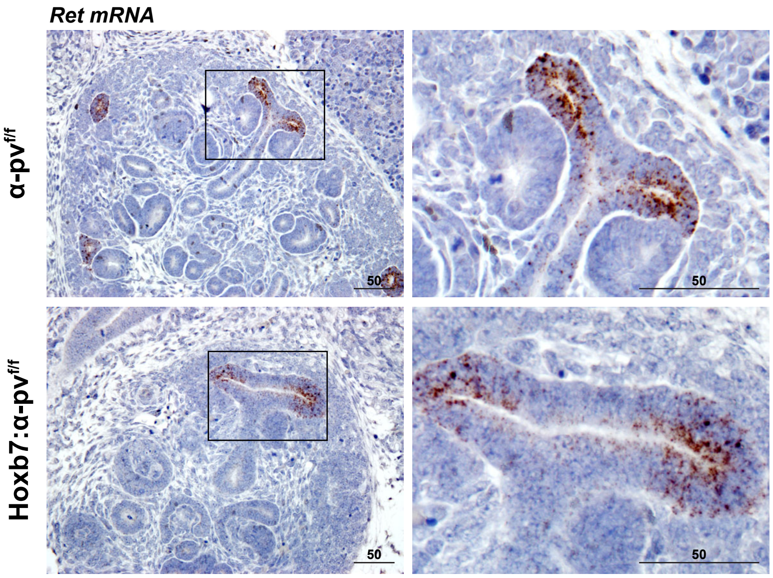

Fig S5

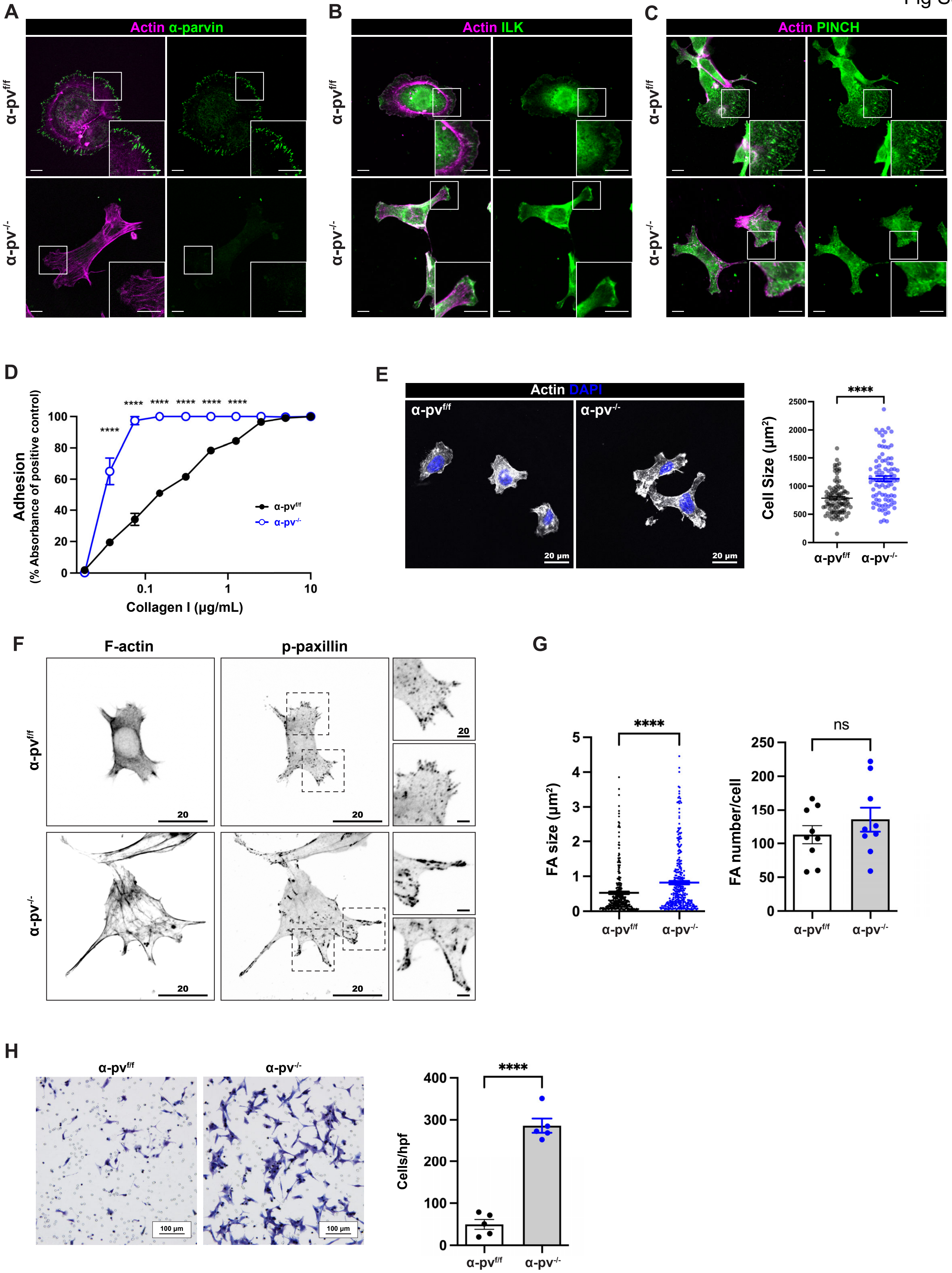
A
Actin α-parvin
α-pv^f/f
α-pv^-/-
B
Actin ILK
C
Actin PINCH
D
Adhesion
(% Absorbance of positive control)
Collagen I (µg/mL)
****
α-pv^f/f
α-pv^-/-
E
Actin DAPI
20 µm
Cell Size (µm²)
****
F
F-actin
p-paxillin
20
G
FA size (µm²)
FA number/cell
ns
H
100 µm
Cells/hpf

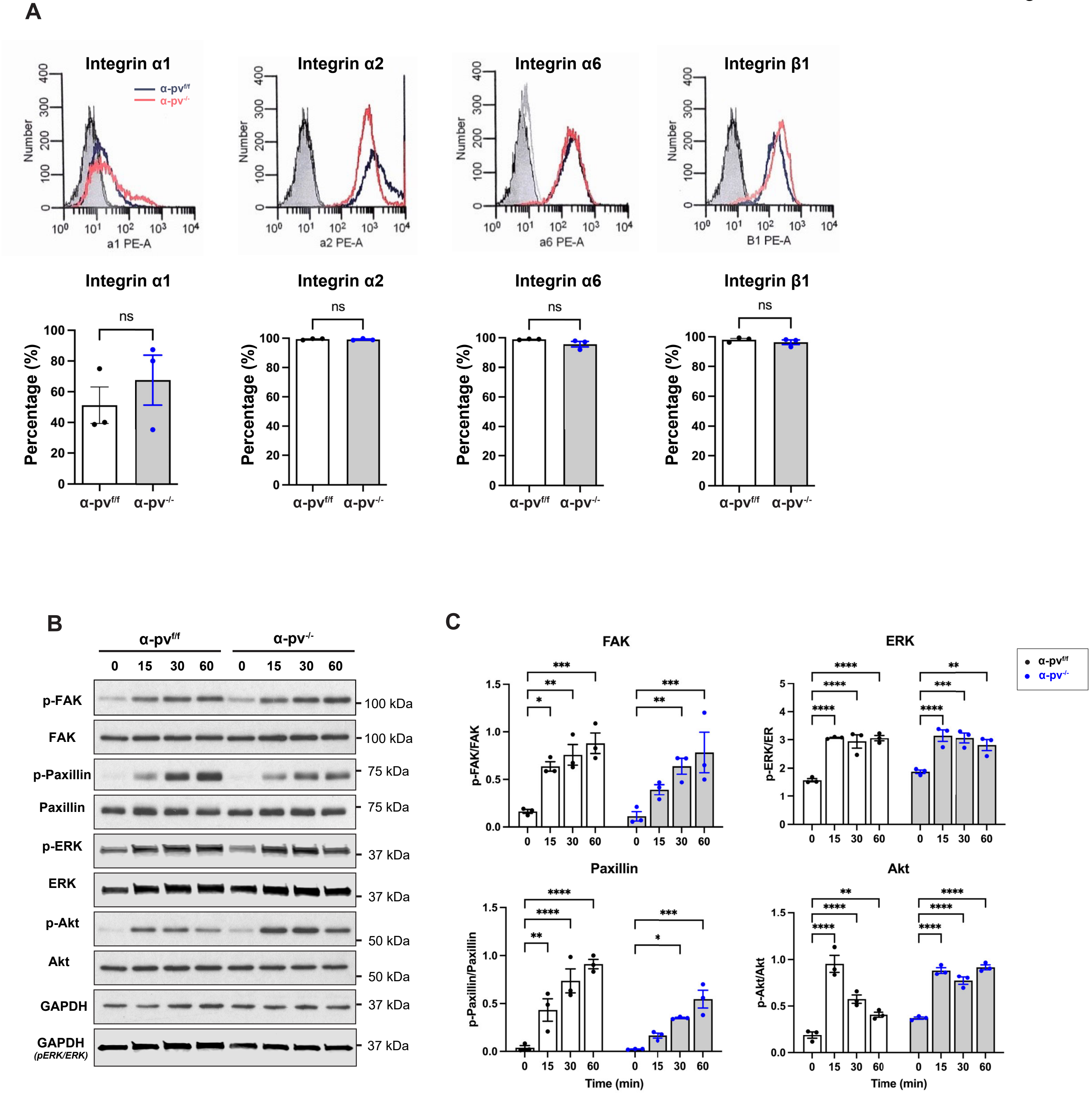
A
Integrin α1
Integrin α2
Integrin α6
Integrin β1
α-pv f/f
α-pv -/-
Number
a1 PE-A
a2 PE-A
a6 PE-A
B1 PE-A
Percentage (%)
ns
α-pv f/f
α-pv -/-
B
0 15 30 60 0 15 30 60
p-FAK
FAK
p-Paxillin
Paxillin
p-ERK
ERK
p-Akt
Akt
GAPDH
GAPDH (pERK/ERK)
100 kDa
75 kDa
37 kDa
50 kDa
C
FAK
ERK
Paxillin
Akt
p-FAK/FAK
p-ERK/ER
p-Paxillin/Paxillin
p-Akt/Akt
Time (min)

Fig S7

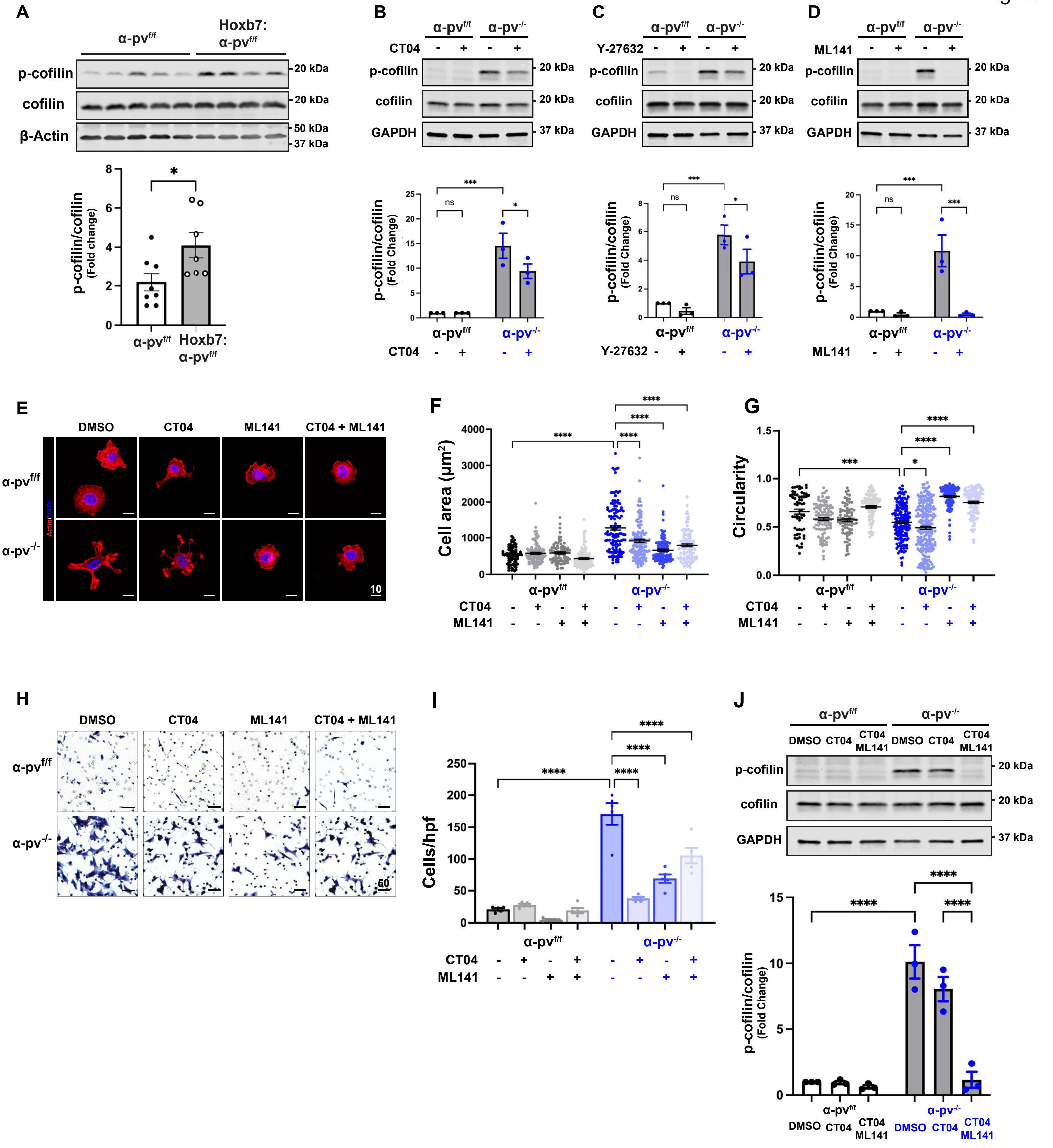

A
α-pv^f/f
Hoxb7: α-pv^f/f
p-cofilin
cofilin
β-Actin
20 kDa
20 kDa
50 kDa
37 kDa
p-cofilin/cofilin (Fold change)
*
α-pv^f/f
Hoxb7: α-pv^f/f
B
α-pv^f/f
α-pv^-/-
CT04
p-cofilin
cofilin
GAPDH
20 kDa
20 kDa
37 kDa
p-cofilin/cofilin (Fold Change)
***
ns
*
C
Y-27632
p-cofilin
cofilin
GAPDH
***
ns
*
D
ML141
p-cofilin
cofilin
GAPDH
***
ns
***
E
DMSO
CT04
ML141
CT04 + ML141
α-pv^f/f
α-pv^-/-
Actin/DAPI
10
F
Cell area (μm²)
****
G
Circularity
****
***
*
H
DMSO
CT04
ML141
CT04 + ML141
α-pv^f/f
α-pv^-/-
50
I
Cells/hpf
****
J
α-pv^f/f
α-pv^-/-
DMSO
CT04
CT04 ML141
p-cofilin
cofilin
GAPDH
20 kDa
20 kDa
37 kDa
p-cofilin/cofilin (Fold Change)
****

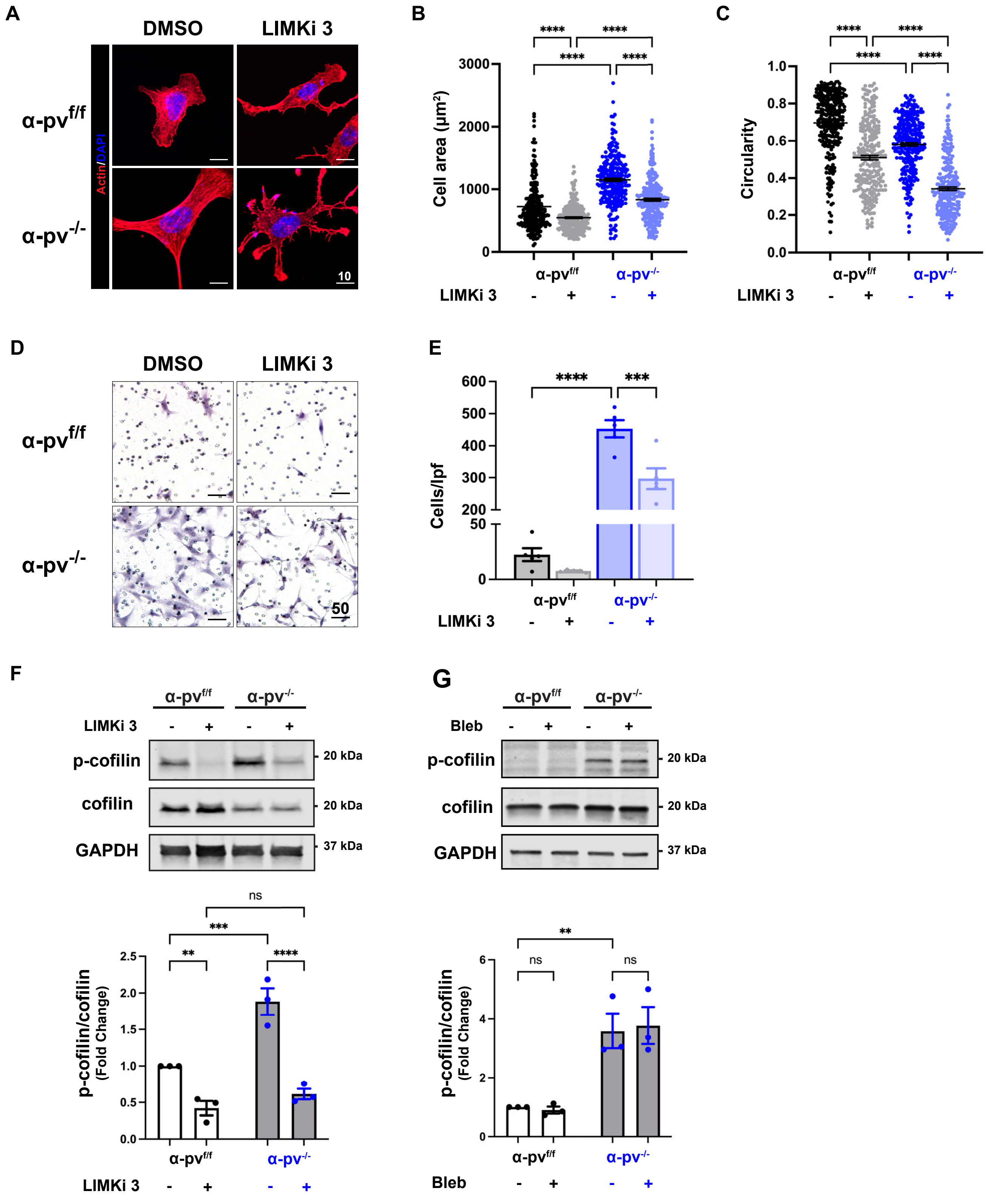
A
DMSO
LIMKi 3
α-pv f/f
α-pv -/-
Actin/DAPI
10
B
Cell area (μm²)
α-pv f/f
α-pv -/-
LIMKi 3
C
Circularity
D
DMSO
LIMKi 3
50
E
Cells/lpf
F
LIMKi 3
p-cofilin
cofilin
GAPDH
20 kDa
37 kDa
p-cofilin/cofilin (Fold Change)
G
Bleb
ns

Fig S9

**A**

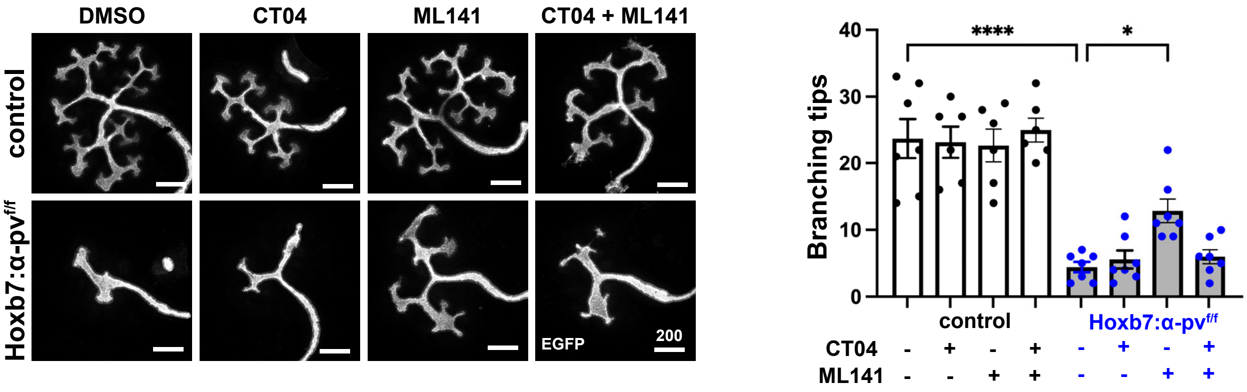


**B**

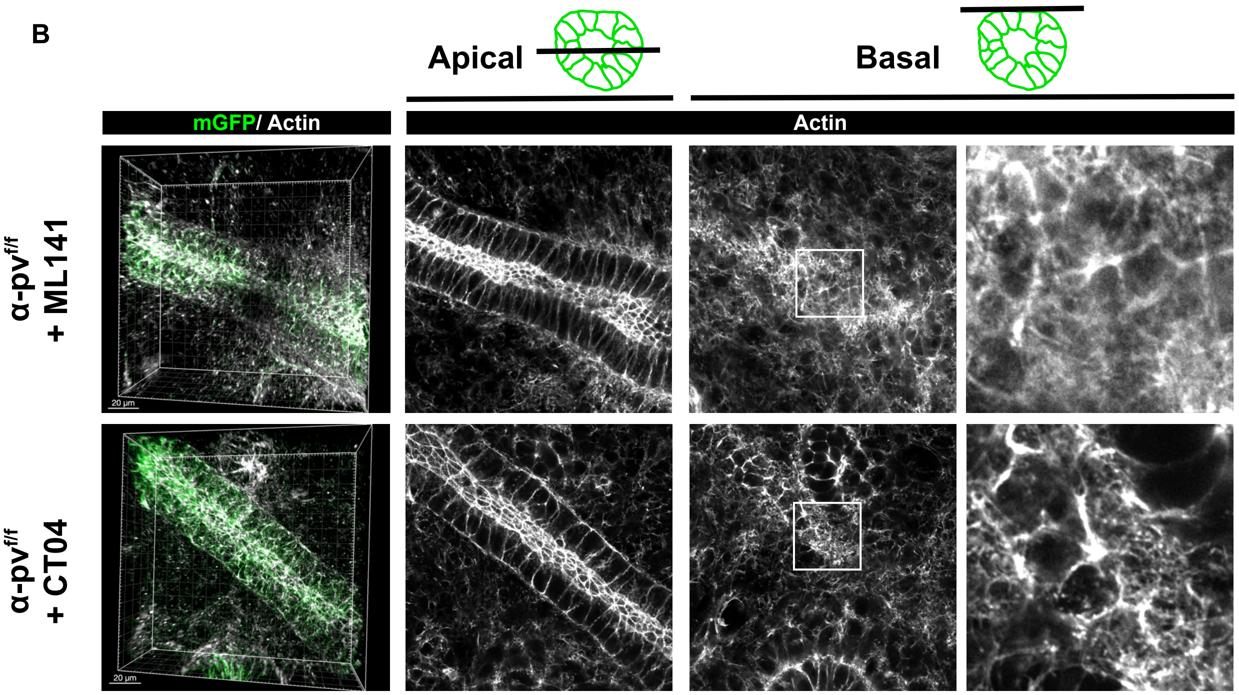


**C**

*ILK$^{f/f}$:α-pv$^{f/f}$ X Hoxb7:α-pv$^{f/+}$*

| | ILK$^{f/+}$:α-pv$^{f/+}$ | Hoxb7: ILK$^{f/+}$:α-pv$^{f/+}$ | ILK$^{f/+}$:α-pv$^{f/f}$ | Hoxb7: ILK$^{f/+}$:α-pv$^{f/f}$ |
|---|---|---|---|---|
| **Expected** | 25% | 25% | 25% | 25% |
| **Observed pups (P14)** | 18.2% (n=4) | 13.6% (n=3) | 63.6% (n=14) | 4.5% (n=1) |

*($p < 0.001$)*

**Supplementary figure 1**

**(A)** E12.5 kidneys were stained for α-parvin (green), E-cadherin (magenta, developing UB), and DAPI (gray) to show that α-parvin is expressed both in the UB trunk and tips of α-$pv^{f/f}$ mice. α-parvin localizes predominantly to the basal side of the UB trunks and localized to the basal-lateral and apical side of the UB tips. α-parvin was deleted in the UB trunks and tips of Hoxb7:α-$pv^{f/f}$ kidneys at E12.5.

**(B)** P0.5 kidneys were stained for α-parvin (green) against WT1 (magenta, podocyte) or LTL (magenta, proximal tubule), and counterstained with DAPI (gray). α-parvin localizes along the glomerular basement membrane (top row), and at the basal side of LTL+ proximal tubules (second row).

**(C)** Hoxb7:α-$pv^{f/f}$ mice are born in the correct Mendelian ratio. Shown in the table are genotypes of offspring obtained from the cross of α-$pv^{f/f}$ and Hoxb7:α-$pv^{f/+}$ mice. The expected numbers of mice were calculated according to the total number of mice born based on the expected Mendelian 1:1:1:1 ratio.

**(D)** Serum BUN was measured in 2-month-old α-$pv^{f/f}$ and Hoxb7:α-$pv^{f/f}$. Shown are mean ± SEM. At least n > 15 mice per group were analyzed. ****$p < 0.0001$; 2-tailed t test

**Supplementary figure 2**

**(A)** Gross anatomy of E18.5 kidneys from α-$pv^{f/f}$ and Hoxb7:α-$pv^{f/f}$ mice. Urogenital systems, including kidneys, ureters, and bladder, were isolated. Hoxb7:α-$pv^{f/f}$ kidneys appeared significantly smaller than α-$pv^{f/f}$ controls, with no evidence of ureteric obstruction.

(**B and C**) Ki67 staining was performed on E18.5 kidneys of α-$pv^{f/f}$ and Hoxb7:α-$pv^{f/f}$ mice where the UB was labelled with cytokeratin (pan) (B, left panel). Ki67 signal within cytokeratin+ UB tubules was filtered using a cytokeratin-based mask (middle panel). Higher magnification is shown in the right panels.. Percentage of ki67-positive CD cells per cross-section were quantified. n = 3 mice per group (G), *$p < 0.05$; 2 tailed t-test.

**Supplementary figure 3**

**(A)** Schematics of cell division angle. An angle of 0° represents a cell division parallel to the longitudinal axis of the tubules. An angle of 90° represents a cell dividing along the apical-basal axis.

**(B)** Examples of cell division angles that were less than (left) or larger than (right) 45º in α-$pv^{f/f}$ (top panels) and Hoxb7:α-$pv^{f/f}$ (bottom panels) tubules. The frames showed rounding of mitotic cells and cytokinesis (magenta dots). Images were taken at 10 min intervals.

**(C)** Cell divisions in both α-$pv^{f/f}$ and Hoxb7:α-$pv^{f/f}$ tubules were randomly oriented when compared to the expected random distribution by the Kolmogorov-Smirnov (KS) test. $p > 0.55$ for both wild-type (n = 50) and mutant (n = 52). Shown are mean ± SEM. 2-tailed t test. n.s. not significant.

**(D)** Stepwise schematics of cell division by luminal mitosis.

**(E)** Cell division event within the UB lumen. Dividing cells (magenta dots) of α-$pv^{f/f}$ (top panels) and Hoxb7:α-$pv^{f/f}$ (bottom panels) UB trunks were imaged at 10 min intervals. The images showed the sequence of the rounding of the premitotic cell, stepwise detachment (arrow) from the basement membrane (cyan line) and cytokinesis. Scale bars represent 10 µm.

**Supplementary figure 4**

RNA scope on E15.5 kidney sections revealed that Ret mRNA was expressed uniformly in cells of the UB tips in both α-pv$^{f/f}$ (top panels) and Hoxb7:α-pv$^{f/f}$ (bottom panels).

**Supplementary figure 5**

**(A-C)** IPP complex localization in the CD cells. CD cells plated on laminin-511 for 4 h were immunostained with α-parvin **(A)**, ILK **(B)** and Pinch **(C)** and co-stained with rhodamine-phalloidin and subjected to confocal microscopy. Scale bar = 10 µm.

**(D)** α-pv$^{-/-}$ and α-pv$^{f/f}$ CD cell adhesion to different concentrations of collagen I at 1 h was measured. Three independent experiments were performed (mean ± SEM), ****p < 0.0001; Two-way ANOVA followed by Tukey's test.

**(E)** α-pv$^{-/-}$ and α-pv$^{f/f}$ CD cells were plated on collagen I and allowed to spread for 1 h, after which they were stained with rhodamine-phalloidin and DAPI. α-pv$^{-/-}$ and α-pv$^{f/f}$ CD cell adhesion area was measured using ImageJ. Shown are mean ± SEM of n > 100 cells per group from a single experiment, which is representative of three consistent independent experiments, ****p < 0.0001; 2 tailed t-test.

**(F-G)** Focal adhesion (FA) formation in the CD cells. CD cells were plated on laminin-511 for 4h and stained with paxillin and α-parvin prior to confocal microscopy. FAs were analyzed in Focal Adhesion Analysis Server (FAAS). FA number was counted and normalized to the cell number or cell area, and FA area were measured **(G)** using Focal Adhesion Analysis Server (FAAS) with values shown in mean ± SEM of 10 individual cells, ****p < 0.0001; 2 tailed t-test.

**(H)** α-pv$^{-/-}$ and α-pv$^{f/f}$ CD cells were plated on transwells coated with collagen I (10 µg/mL) and allowed to transmigrate through 8 um pores for 4 h. Cells were counted in 5 high-power fields per group. Five independent experiments were performed (mean ± SEM), ****p < 0.0001; 2 tailed t-test.

**Supplementary figure 6**

**(A)** Surface expression of integrin β1, α1, α2, α3, and α6 subunits was analyzed by FACS in α-pv$^{f/f}$ and α-pv$^{-/-}$ CD cells by using R-phycoerythrin conjugated secondary antibodies.

**(B and C) α-pv**$^{f/ff}$ and **α-pv**$^{-/-}$ CD cells were plated on laminin-511 for 0, 15, 30 and 60 min and immunoblotted for phosphorylated and total FAK, Paxillin, ERK and Akt. GAPDH was blotted to verify equal protein loading (A). Three independent experiments were quantified using densitometry and shown as individual values and mean ± SEM. p-ERK/ERK were blotted on a different membrane with a separate loading control as indicated (B). *p < 0.05, **p < 0.01, *** p < 0.001, ****p < 0.0001; Two-way ANOVA followed by Tukey's test.

**Supplementary figure 7**

**(A)** Papillary lysates from α-pv$^{f/f}$ and Hoxb7:α-pv$^{f/f}$ kidneys were subject to immunoblotting for phosphorylated and total cofilin. Quantification of at least seven mice per group is shown in the lower panel (mean ± SEM). *p < 0.05; 2 tailed t-test.

**(B-D)** α-pv$^{f/f}$ and α-pv$^{-/-}$ CD cells were treated with CT04 (B), Y-27632 (C), ML141 (D), and immunoblotted for phosphorylated and total cofilin. Three experiments were quantified using densitometry and shown as individual values and mean ± SEM in the lower panel. *p < 0.05, **p < 0.001, ***p < 0.0001; Two-way ANOVA followed by Tukey's test.

**(E, F and G)** α-pv$^{f/f}$ and α-pv$^{-/-}$ CD cells were treated with the Rho inhibitor CT04 (2 µg/mL), the Cdc42 inhibitor (10 µM ) or a combination of Rho and Cdc42 inhibitors (CT04 2 µg/mL and ML141 10 µM) and allowed to spread on Laminin-511 for 1h. Representative images of rhodamine-phalloidin–stained cells (E) are shown. Spreading area (F) and circularity (G) were quantified using FIJI. Shown are mean ± SEM of n > 100 cells per group from a single experiment, which is representative of three consistent independent experiments, ****p < 0.0001; Two-way ANOVA followed by Tukey's test.

**(H and I)** α-pv$^{f/f}$ and α-pv$^{-/-}$ CD cells were treated with the Rho inhibitor CT04 (2 µg/mL), the Cdc42 inhibitor (10 µM ) or a combination of Rho and Cdc42 inhibitors (CT04 2 µg/mL and ML141 10 µM) and subjected to a 4h transwell migration assay. Representative images of crystal violet-stained cells (H) are shown. Migrated cells (I) were counted manually in FIJI. Five independent experiments were performed (mean ± SEM) with at least 5 low-power fields per group counted. ****p < 0.0001; Two-way ANOVA followed by Tukey's test.

**(J)** α-pv$^{f/f}$ and α-pv$^{-/-}$ CD cells were treated with Rho inhibitor CT04 (2 µg/mL) or a combination of Rho and Cdc42 inhibitors (CT04 2 µg/mL and ML141 10 µM) and immunoblotted for phosphorylated and total cofilin. Three experiments were quantified using densitometry and shown as individual values and mean ± SEM in the lower panel. *p < 0.05, **p < 0.001, ***p < 0.0001; Two-way ANOVA followed by Tukey's test.

**Supplementary figure 8**
**(A-C)** α-pv$^{f/f}$ and α-pv$^{-/-}$ CD cells were treated with the LIMK inhibitor LIMKi 3 (10 µM) and allowed to spread on Laminin-511 for 1h. Representative images of rhodamine-phalloidin–stained cells (A) are shown. Spreading area (B) and circularity (C) were quantified using FIJI. Shown are mean ± SEM of n > 100 cells per group from a single experiment, which is representative of three consistent independent experiments, ****p < 0.0001; Two-way ANOVA followed by Tukey's test.

**(D and E)** α-pv$^{f/f}$ and α-pv$^{-/-}$ CD cells were treated with LIMK inhibitor LIMKi 3 (10 µM) and subjected to a 4h transwell migration assay. Representative images of crystal violet-stained cells (D) are shown. Migrated cells (E) were counted manually in FIJI. Five independent experiments were performed (mean ± SEM) with at least 5 low-power fields per group counted. ***p < 0.001, ****p < 0.0001; Two-way ANOVA followed by Tukey's test.

**(F)** α-pv$^{f/f}$ and α-pv$^{-/-}$ CD cells were treated with LIMK inhibitor LIMKi 3 (10 µM) and immunoblotted for phosphorylated and total cofilin. Three experiments were quantified using densitometry and shown as individual values and mean ± SEM in the lower panel. **p < 0.01, ***p < 0.001, ****p < 0.0001; Two-way ANOVA followed by Tukey's test.

**(G)** α-pv$^{f/f}$ and α-pv$^{-/-}$ CD cells were treated with Blebbistatin (10 µM) and immunoblotted for phosphorylated and total cofilin. Three experiments were quantified using densitometry and shown as individual values and mean ± SEM in the lower panel. **p < 0.01; Two-way ANOVA followed by Tukey's test.

**Supplementary figure 9**

**(A)** E11.5 control (control kidneys were Hoxb7:mTmG$^{f/f}$ unless otherwise noted.) and Hoxb7:α-pv$^{f/f}$ kidneys were isolated and grown ex vivo in the presence of a combined Rho and Cdc42 inhibitor (CT04 2 μg/mL, ML141 10μM) for 24h and imaged by confocal microscopy. Representative images demonstrate a partial rescue of the branching defect in the **Hoxb7:α-pv$^{f/f}$** kidneys.. Branching points were counted in ImageJ with data (mean ± SEM) shown on the right panel. A minimum of 7 embryonic mice were included per group. Each treated kidney was compared with its respective untreated contralateral kidney. The experiments were conducted in three independent sets. **p < 0.001; paired 2 tailed t-test.

**(B)** E11.5 kidneys isolated from control mice were cultured ex vivo with a combined Rho and Cdc42 inhibitor (CT04 2 μg/mL, ML141 10μM) for 24 hours and imaged by confocal microscopy. UB trunks were visualized by GFP staining, and actin filaments were visualized with rhodamine-conjugated phalloidin. Panels from left to right show: the 3D reconstruction of the UB trunk, apical section, basal section, and the higher magnification of the basal actin.

(C) A double floxed ILK$^{f/f}$:α-pv$^{f/f}$ line was crossed with Hoxb7:α-pv$^{f/+}$ mice to generate offspring heterozygous for ILK in an α-parvin-null background (Hoxb7:ILK$^{f/+}$:α-pv$^{f/f}$). Based on Mendelian ratios, 25% of pups were expected to carry the Hoxb7:α-pv$^{f/f}$ genotype; however, genotyping at P14 revealed significantly fewer Hoxb7:ILK$^{f/+}$:α-pv$^{f/f}$ mice and an excess of Cre-negative littermates (ILK$^{f/+}$:α-pv$^{f/f}$), deviating from the expected distribution ($\chi^2 = 18.37$, $p < 0.001$).

**Supplementary Movies**

**Movies 1 and 2.** Time-lapse fluorescence microscopy showing the 3D rendering of a branching tip from an E11.5 mTmG transgenic mouse.

**Movies 3 and 4.** Orthogonal view time-lapse microscopy of the branching tip from an E11.5 mTmG transgenic mouse. Images showing x-y, y-z and x-z plane.

**Movies 5 and 6.** Time-lapse microscopy reveals cell rearrangements in rosette-like clusters in the ureteric bud of mTmG transgenic mice. Cells within these clusters were manually segmented and colored (orange), tracked over a 6-hour period at 10-minute intervals.

**Movies 7 and 8.** Simulations showing divergency map flow arrows.

**Movies 9 and 10.** Simulations showing similarity maps.
